# TOWARD A CONNECTIVITY GRADIENT-BASED FRAMEWORK FOR REPRODUCIBLE BIOMARKER DISCOVERY

**DOI:** 10.1101/2020.04.15.043315

**Authors:** Seok-Jun Hong, Ting Xu, Aki Nikolaidis, Jonathan Smallwood, Daniel S. Margulies, Boris Bernhardt, Joshua Vogelstein, Michael P. Milham

**Author notes:** These authors contributed equally (Seok-Jun Hong, Ting Xu). **Corresponding authors**, Michael P. Milham, MD, PhD, Center for the Developing Brain, Child Mind Institute, Center for Biomedical Imaging and Neuromodulation, Nathan Kline Institute, NY, USA, 101 East 56th Street, New York, NY 10022, Seok-Jun Hong, PhD, Center for the Developing Brain, Child Mind Institute, NY, USA, Center for Neuroscience Imaging Research, Institute for Basic Science, Department of Biomedical Engineering, Sungkyunkwan University, Suwon, South Korea.

## Abstract

Despite myriad demonstrations of feasibility, the high dimensionality of fMRI data remains a critical barrier to its utility for reproducible biomarker discovery. Recent studies applying dimensionality reduction techniques to resting-state fMRI (R-fMRI) have unveiled neurocognitively meaningful connectivity gradients that are present in both human and primate brains, and appear to differ meaningfully among individuals and clinical populations. Here, we provide a critical assessment of the suitability of connectivity gradients for biomarker discovery. Using the Human Connectome Project (discovery subsample=209; two replication subsamples= 209×2) and the Midnight scan club (n=9), we tested the following key biomarker traits – reliability, reproducibility and predictive validity – of functional gradients. In doing so, we systematically assessed the effects of three analytical settings, including *i*) dimensionality reduction algorithms (*i.e*., linear *vs*. non-linear methods), *ii*) input data types (*i.e*., raw time series, [un-]thresholded functional connectivity), and *iii*) amount of the data (R-fMRI time-series lengths). We found that the reproducibility of functional gradients across algorithms and subsamples is generally higher for those explaining more variances of whole-brain connectivity data, as well as those having higher reliability. Notably, among different analytical settings, a linear dimensionality reduction (principal component analysis in our study), more conservatively thresholded functional connectivity (*e.g*., 95-97%) and longer time-series data (at least ≥20mins) was found to be preferential conditions to obtain higher reliability. Those gradients with higher reliability were able to predict unseen phenotypic scores with a higher accuracy, highlighting reliability as a critical prerequisite for validity. Importantly, prediction accuracy with connectivity gradients exceeded that observed with more traditional edge-based connectivity measures, suggesting the added value of a low-dimensional gradient approach. Finally, the present work highlights the importance and benefits of systematically exploring the parameter space for new imaging methods before widespread deployment.

**Highlights:**

- There is a growing need to identify benchmark parameters in advancing functional connectivity gradients into a reliable biomarker.
- Here, we explored multidimensional parameter space in calculating functional gradients to improve their reproducibility, reliability and predictive validity.
- We demonstrated that more reproducible and reliable gradient markers tend to have higher predictive power for unseen phenotypic scores across various cognitive domains.
- We showed that the low-dimensional connectivity gradient approach could outperform raw edge-based analyses in terms of predicting phenotypic scores.
- We highlight the necessity of optimizing parameters for new imaging methods before their widespread deployment.

## Introduction

Inspired by the resurgence of dysconnectivity models in neuropsychiatric disorders over the past decade (Catani and Ffytche, 2005; van den Heuvel and Sporns, 2019), research targeting brain connectivity has become a central focus of imaging-based biomarker discovery in clinical neuroscience (Castellanos et al., 2013; Di Martino et al., 2014). Early neuroimaging studies have targeted specific connections/networks of interest, often motivated by prior neuropsychological models of brain dysfunction (Garrity et al., 2007; Greicius et al., 2007; Roalf and Gur, 2017). However, as alternative conceptual views emphasized the importance of understanding whole brain network topology, more recent work has characterized the system-level principles of brain organization (Menon, 2011; van den Heuvel and Sporns, 2019). Resting state fMRI (R-fMRI) has been particularly useful in these efforts (Castellanos et al., 2013), as it made major findings in functional neuroimaging possible, including discovery of canonical functional brain networks (Craddock et al., 2012; Yeo et al., 2011), identification of areal boundaries based on connectivity profiles (Cohen et al., 2008; Wig et al., 2014) and characterization of graph-theoretical properties for network topology (*e.g*., small-worldness, centrality, rich-club) (Bullmore and Sporns, 2009; Van Den Heuvel and Sporns, 2011). Recognizing the high dimensionality of functional connectivity data, however, emerging efforts have highlighted the need for identifying summary metrics that can distill complex whole-brain connectivity data into more parsimonious sets of organizing principles. Toward this goal, a framework has been introduced to reduce such complexity into a set of dimensions describing the ‘connectivity space’ of the brain (Haak et al., 2018; Langs et al., 2016; Margulies et al., 2016; Mars et al., 2018a, 2018b; Vos de Wael et al., 2020). Despite the value of these approaches, there is currently a lack of consensus on which method is the most applicable to develop an effective imaging biomarker.

In the present work, we sought to address this missing gap and advance connectome-based biomarker discovery by systematically assessing the reliability and predictive validity of lowdimensional representations of whole-brain functional connectivity. By applying a dimensionality reduction algorithm to whole-brain functional connectivity data, this approach has effectively unveiled multiple primary axes – referred as ‘gradients’ (or more formally ‘manifolds’ in the case of non-linear dimensionality reduction). These gradients describe smooth transitions of functional connectivity patterns along the cortical surface. Indeed, a recent study using this technique (Margulies et al., 2016) revealed several important principal systems of large-scale cortical hierarchy such as the transition from sensory to transmodal areas, the segregation of the primary sensory/motor systems (Hilgetag and Goulas, 2020; Mesulam, 1998) and the pattern spanning across an intrinsic to task-positive or multiple-demand network (Duncan, 2010; Fox et al., 2005). A low-dimensional representation of functional connectivity, therefore, provides a unified perspective to efficiently explain cognitively plausible macro-scale mechanisms of the functional brain, which has led to these approaches to recently gain increasing attention in the neuroimaging community.

Prior work has examined connectivity gradients and their relations using a variety of methods and analytic choices. One approach is to focus on specific regions, such as the striatum, sensorimotor areas and mesial temporal lobe structures including the entorhinal cortex and hippocampus (Haak et al., 2018; Marquand et al., 2017; Navarro Schröder et al., 2015; Przeździk et al., 2019; Vos de Wael et al., 2018). These functional connectivity profiles reveal ordered changes along the primary axes, which often effectively recapitulate underlying anatomical connectivity transition across the brain areas (Marquand et al., 2017). Other studies have also examined how the connectivity gradients are related to ongoing cognition. These have revealed that the principal gradient is linked to patterns where the cognition is guided by information from memory rather than sensory input (Murphy et al., 2019, 2018) and contributes to detailed representations of task-relevant functional states (Sormaz et al., 2018). One of the most important uses for the gradient approach, however, is to identify biomarkers for clinical samples. Hong, *et al*. demonstrated autism-related decreases in the separation of brain networks along the hierarchy that describes transitions from unimodal to transmodal cortices, and demonstrated that these changes were predictive of social and behavioral difficulties that these individuals reported (Hong et al., 2019). While these studies supported feasibility and potential clinical utility for the use of low-dimensional connectome representations, key issues remain. Importantly, a broad catalogue of relevant parameters and algorithms results in a high level of analytic degrees of freedom. As such, there is a growing need to identify benchmark parameters that provide an efficient and reliable parameter space for connectivity gradients. This call is particularly urgent, because the software packages to calculate gradients have been increasingly more available (*e.g*., BrainSpace, https://brainspace.readthedocs.io/en/latest/, Vos de Wael et al., 2020; congrads, https://github.com/koenhaak/congrads, Haak et al., 2018). These efforts will likely accelerate the pace of future studies, and without benchmark parameters, they will yield the findings only based on suboptimal analytic settings.

Here, we provided a quantitative assessment of gradient-based measures for usage in biomarker discovery studies based on multiple openly shared large-sample datasets. Central to this goal, we have explored the parameter options for algorithmic decisions, including: *i*) gradient extraction method (*i.e*., principal component analysis [PCA], Jolliffe, 2011; diffusion map embedding [DE], Coifman et al., 2005; Laplacian Eigenmaps [LE], Belkin and Niyogi, 2003), *ii*) data representation (*i.e*., timeseries, un-thresholded and thresholded functional connectivity matrices), *iii*) amount of functional imaging data included (5-50 minutes), and *iv*) their impact on univariate and multivariate indices of reliability (*i.e*., intraclass correlation coefficient, Shrout and Fleiss, 1979; discriminability, Bridgeford et al., 2020). To ensure that these measures have practical value, we also explored the ability of gradients to predict various phenotypic measures, including cognition, personality, and psychiatric symptoms. Finally, we assessed between-algorithm and -sample reproducibility, critical aspects of valid and robust imaging biomarkers.

## Methods

### General analytic flow

**Table 1** outlines a four-fold analytic strategy of the current study. *Analysis-1* examined raw functional gradient profiles from three different dimensionality reduction algorithms (*i.e*., PCA, DE, LE). We assessed the similarity of gradients produced by the different algorithms, as well as their reproducibility in a non-overlapping sample. *Analysis-2* evaluated the reliability of each gradient using two established metrics (*i.e*., ICC [univariate] and discriminability [multivariate], Bridgeford et al., 2020), and compared them across different algorithms and input data representation (*i.e*. time-series *vs*. functional connectivity with different thresholds). *Analysis-3* evaluated the effect of R-fMRI time-series length (5 to 50mins) on the reliability. Finally, *Analysis-4* assessed prediction accuracy of each gradient for various behavioral and cognitive outcomes, systematically varying the algorithm and input type as well as R-fMRI time-series length.

**TABLE 1.** Summary of Objectives, Materials, Methods and Key Findings.

| Aims | Analytic methods | Datasets | Key findings |
| --- | --- | --- | --- |
| **Analysis-1**<br>To assess raw functional gradients and their reproducibility | 1. <u>Profiling of gradients</u><br>a. Using different algorithms (*i.e.* PCA, DE, LE)<br>b. Using different input data types (*i.e.* time-series, un-thresholded FC, and thresholded: 5, 25, 50, 75, 90, 95, 96, 97, 98, 99%)<br>c. Estimating the variance explained by each component across the algorithms<br>2. <u>Reproducibility test</u>: Correlating extracted gradients<br>a. Between the three algorithms<br>b. Between discovery and replication datasets | Concatenated two R-fMRI time-series (=2×15=30mins) in both two sessions (total: two 30mins time-series for test and retest) in 209 subjects (discovery & replication) | 1. First four gradients are similar across algorithms (**FIGURE 1A, C**).<br>2. Discrepant patterns from the 5<sup>th</sup> gradient can be aligned using a Procrustes linear transformation (**FIGURE 1C**).<br>3. PCA and LE show the most similar gradient patterns, whereas PCA and DE reveal the least similar patterns (**FIGURE 1C**).<br>4. The discovery-replication reproducibility is generally high ($r>0.8$) until the 6<sup>th</sup> gradient (even if they are in raw order), and further increased after a gradient matching process (**FIGURE 1C**).<br>5. As FC threshold is increased, variance explained becomes less dominated by first primary gradients, regardless of algorithm used (**FIGURE 1B**). |
| **Analysis-2**<br>To compare the reliability of gradients across the algorithms and input data types | 1. <u>Profiling of intraclass correlation coefficient (ICC)</u><br>a. Calculating ICC at each gradient across the algorithms and across different input data types<br>b. Counting # of gradients with ICC>0.5 to compare between algorithms and input data types<br>c. Detecting the most reliable gradient among those from different parameter setting and visualizing its whole-brain pattern<br>2. <u>ICC comparison between the algorithms</u><br>a. Constructing a similarity matrix of ICC between the algorithms<br>b. Assessing between- and within-subject variability<br>3. <u>Discriminability comparison</u> at each gradient across the algorithms and input data types. | Concatenated two R-fMRI time-series (=2×15=30mins) in both two sessions (total: two 30mins time-series for test and retest) in 209 subjects (discovery & replication) | Regardless of the algorithm used, gradient reliability:<br>1. is positively associated with the variance explained (*i.e.*, gradient order, **FIGURE 2A**).<br>2. increases with higher FC thresholds (**FIGURE 2A**).<br>3. is maximized in the 95-97% range of FC threshold.<br>4. is maximum in the default mode, fronto-parietal and dorsal attention networks, mainly due to a high between-subject variability (**FIGURE 2B**).<br>5. is highly reproducible (**SUPPLEMENTARY FIGURE 3**).<br>Between the different algorithms,<br>1. PCA generally shows higher ICC (**FIGURE 2A**).<br>2. However, DE shows higher reliability in the range of minor gradients (from about 40<sup>th</sup> gradient order; **FIGURE 2A**).<br>3. PCA and DE show high discriminability (nearly perfect values when using FC matrices with a high threshold (**FIGURE 2C**)). |
| **Analysis-3**<br>To assess effects of R-fMRI time-series length on the reliability of functional gradients | 1. <u>Evaluation of ICC change as a function of R-fMRI time-series length across the three algorithms</u><br>a. Calculating ICC of the gradient from 95%-thresholded FC across 5, 10, 20, 30, 40, 50 mins of time-series data<br>b. Categorizing the ICC into the 'fair', 'moderate', 'substantial' and 'almost perfect' intervals and assessing the proportions across different scan times.<br>c. Assessing the time-series length-dependent ICC changes across different functional networks.<br>2. <u>Evaluation of discriminability as a function of time-series length across the three algorithms</u> | Midnight Scan Club R-fMRI time-series of 9 subjects (each having ten 30mins of data=5 hours of R-fMRI data) | 1. As the time-series length gets longer, reliability is monotonously increased over the whole brain without distinct patterns across the functional subnetworks (**FIGURE 3A**)<br>2. PCA shows higher reliability than other algorithms regardless of scan duration in all categories (*i.e.*, moderate, substantial and almost perfect, **FIGURE 3A** middle panels)<br>3. In general, from-30-to-40 mins of R-fMRI data change makes the largest jump of ICC increase (**FIGURE 3A** bottom panels).<br>4. The gradients explaining more variance reveal higher discriminability compared to the minor gradients (**FIGURE 3B**). |
| **Analysis-4**<br>To evaluate prediction power of gradients based on various combinations of parameter setting, and to relate this accuracy to the reliability | 1. <u>Testing a prediction accuracy</u><br>a. Reducing the dimensionality of phenotypic data using an exploratory factor analysis (EFA)<br>b. Performing a prediction test on the factor scores and entire phenotypic scores<br>2. <u>Relating a prediction accuracy to reliability of gradients</u><br>3. <u>Assessing prediction accuracy across different algorithms and input data types</u><br>a. Comparing between algorithms and between input data types<br>b. Checking the spatial patterns of contributing features across the whole brain<br>4. <u>Assessing the prediction accuracy across different R-fMRI time-series lengths</u><br>5. <u>Comparing the prediction accuracy with connectome-based predictive modeling</u> | Concatenated two R-fMRI time-series (=2×15=30mins) in both two sessions (total: two 30mins time-series for test and retest) in 209 subjects and 65 phenotypic scores in 3 datasets of 209 subjects (training, test1 and test2) | 1. EFA identified 3 factors from entire phenotypic scores, representing, externalizing and externalizing symptoms, and general cognitive function (**SUPPLEMENTARY FIGURE 1**)<br>2. General cognitive function was predicted in many more gradients (especially for the components explaining larger data variance) compared to internalizing and externalizing factor scores (**FIGURE 4A, B**). The findings were highly reproducible in the test-2 dataset.<br>3. Averaged prediction accuracy was significantly associated gradient reliability (both ICC and discriminability (**FIGURE 4A**)).<br>4. Across all choices of algorithms and thresholds, prediction accuracy was maximized with PCA using a 95-6% threshold (**FIGURE 5**)<br>5. The brain regions with higher reliability tend to show greatest feature contributions to CCA-based prediction (**FIGURE 5**)<br>6. The gradients outperform connectome-based predictive modeling in terms of phenotypic score prediction (**FIGURE 6**)<br>7. 5-10 minutes of data are insufficient to reliably predict phenotypic scores. 20 minutes may be a reasonable compromise between clinical feasibility and biomarker optimization (**FIGURE 7**). |

### Data and code availability statement

In performing these analyses, we used Matlab 2017b as our main computing platform. Specifically, PCA was calculated based on pca.m implemented in the ‘*Statistics and Machine Learning Toolbox*’, while DE and LE calculation, as well as gradient alignment, were done using the BrainSpace toolbox (https://brainspace.readthedocs.io/en/latest/, Vos de Wael et al., 2020). ICC was computed using a function (IPN_icc.m) from the Connectome Computation System (Xu et al., 2015, zuoxinian/CCS: Connectome Computation System), which was implemented based on the established approach (Shrout and Fleiss, 1979). Finally, discriminability was calculated by compute_mnr.m in neurodata/discriminability.

The data analyzed in this study were all downloaded from two open source data repositories: the Human Connectome Project (HCP, Glasser et al., 2013) and the Midnight Scan Club (MSC, Gordon et al., 2017). All the codes and the list of the subjects used in this study are released in ChildMindInstitute/GradientBiomarker.

### Subjects and imaging-behavior data

In HCP, we selected three independent sets to demonstrate the reproducibility of our findings: *i*) discovery (or training set for prediction analyses; 209 subjects; age: 27.7±3.7ys, 106 males), *ii*) replication-1 (or test-1 set in prediction analyses; 209 subjects, age: 28.4±4.0ys, 106 males) and *iii*) replication-2 (or test-2 set in prediction analyses; another 209 subjects; age: 29.3±3.4ys, 74 males). Notably, in discovery and replication-1 datasets, we selected the subjects based on their genetic unrelatedness (*i.e*. no overlapped family member) to rule out inflated reproducibility or prediction accuracy in our findings. The replication-2 was a subset of remaining subjects whose family members overlapped with those of either discovery or replication-1 subjects. From these cases, we included R-fMRI and phenotypic scores, which were analyzed in *Analysis-1, −2* and *−4* (see **Table 1**). On the other hand, in *Analysis-3*, to assess different R-fMRI time-series effects based on more densely sampled individual fMRI data, we chose the MSC data that has 9 subjects (age: 29.1±3.3ys, 5 males), each having 10 sessions of 30-mins R-fMRI data.

1. <u>HCP</u>: The description about R-fMRI and behavioral data of HCP is detailed elsewhere (Barch et al., 2013; Glasser et al., 2013; Smith et al., 2013). Briefly, R-fMRI data was acquired from a 3T Siemens connectome-Skyra scanner using a gradient-echo EPI sequence (TE=33.1ms, TR=720ms, flip angle = 52°, 2.0mm isotropic voxels, 72 slices, multiband factor of 8) at each individual. The data was obtained through two sessions, each of which ran two R-fMRI scans (each approximately 15 minutes). The two R-fMRI scans were acquired for different phase encoding directions (left-right [LR] and right-left [RL] scans), and thus two sessions provided 4 R-fMRI datasets in total ([LR1, RL1, LR2, RL2] × ~15mins = ~60mins [4×1200volumes]). The HCP also provided a rich array of phenotypic scores for each participant (Barch et al., 2013). Among the available phenotypic scores, we selected 6 major cognitive and psychiatric domains (*i.e*., alertness, cognition, emotion, personality, sensory and psychiatric and life functions; 65 different individual scores in total), and associated the raw scores and their factor scores (see **Supplementary Figure 1** for specific items and factors) to functional gradients.
2. <u>MSC</u>: Acquisition details are specified in the original data descriptor paper (Gordon et al., 2017). We included R-fMRI datasets from 9 subjects who underwent 30 minutes of a gradient-echo EPI scan (TE = 27ms, TR = 2.2s, flip angle = 90°, 4.0mm isotropic voxels, 36 slices) over ten subsequent days (30mins×10=5 hours [818×10 volumes] in total). We did not include behavioral data from MSC.

### fMRI preprocessing and parcellation

1. <u>HCP</u>: We used R-fMRI data that already underwent HCP’s minimal preprocessing pipeline (Glasser et al., 2013). No slice timing correction was performed, spatial preprocessing was applied, and structured artefacts were removed using ICA+FIX (independent component analysis followed by FMRIB’s ICA-based X-noiseifier, Salimi-Khorshidi et al., 2014), which is known for its ability to remove >99% of the artefactual components from a given dataset. Cleaned R-fMRI data was represented as a time-series of grayordinates (*i.e*., a combination of cortical surface vertices and subcortical standard-space voxels) and bandpass filtered at 0.008-0.08 Hz. We discarded the first 10 volumes (7.2sec) to allow the magnetization to stabilize to a steady state (=1190 volumes). We downsampled the original 32k time-series into those with 10k vertices to accelerate further preprocessing steps. To generate test-retest datasets while reducing potential session-related batch effects, we concatenated normalized signals (*i.e*. z-score) of LR1 and RL2 (see a previous ‘*1. HCP*’ section for acronyms) images (test; 2380 volumes), and also those of RL1 and LR2 images (retest; 2380 volumes), each yielding ~30mins of time-series across the individuals. To reduce computational cost for gradient calculation, we averaged vertex-wise time-series into a larger-size of regions of interest across the whole brain using a Schaefer parcellation (Schaefer et al., 2018). While the original parcellation has 1000 ROIs on the 32k cortical surface, 2 ROIs were discarded during the 10k downsampling due to their small parcel size. This provided two sets of 998×2380 time-series matrices (test and retest) at each individual, which became the main inputs for following gradient calculation.
2. <u>MSC</u>: We have used the data that were already preprocessed by the MSC imaging pipeline (Gordon et al., 2017). The steps included slice-timing correction, frame-to-frame alignment for head motion correction, intensity normalization and distortion correction. Preprocessed fMRI data was registered to anatomical images and sampled onto the vertices of extracted 32k cortical surfaces (fs_LR_32k). Again, we downsampled the original 32k grayordinates time-series into those with 10k vertices and averaged the time-series signals into 998 ROIs of the Shaefer’s parcellation map. This provided 10 sets of 998×818 time-series matrices at each individual, which were used to assess the effects of the amount of data in following analyses.

### Analysis-1. Gradient extraction, alignment, and reproducibility tests

While current literature for dimensionality reduction highlights the strengths of many conceptually distinct methods, here we focused on three primary ones, including the most representative and simple linear algorithm (*i.e*. principal component analysis) and two nonlinear manifold learning methods that have been previously used in the neuroimaging field (*i.e*., diffusion embedding map, Coifman and Lafon, 2006; Laplacian Eigenmaps, Belkin and Niyogi, 2003). Another factor that may affect the gradient calculation is the type of signals that are entered into the algorithm. Centered at row-wise 90% thresholding (*i.e*., a threshold leaving only top 10% of the strongest functional connectivity at each brain area; a method proposed in previous studies, Margulies et al., 2016; Yeo et al., 2011), we have applied a systematically varied threshold from 0, 25, 50, 75, 90, 95, 96, 97, 98 and 99% on a functional connectivity matrix to see which threshold would lead to the most reliable gradients across the whole brain. We also tested the effect of time-series signals without constructing a connectivity matrix, aiming to assess if raw time-series may already show high reliability. The combination of these parameters and algorithms yielded 3 (# of different algorithms) × 11 (different input data representation) pairs of functional gradient results.

### 1. Gradient extraction

a. <u>Principal Component Analysis (PCA)</u>: PCA was performed using singular vector decomposition (SVD: X=USV^**T**^; X: time series or a [un]thresholded functional connectivity matrix, U: left-singular vectors, S: a diagonal matrix of singular values, V: right-singular vectors). In an SVD setting, the columns of V represent principal directions (axes) of cortical points for which distance is determined by their functional connectivity. The columns of U×S are principal components or scores of brain areas projected onto those identified principal axes (so called “functional gradient”). S is related to the eigenvalues of covariance matrix via *λ_i_*=S_i_^2^/(n-1), which can later be used to estimate variances explained by each principal component. We applied SVD to both time series (998×2380) and functional connectivity matrices (998×998) of each individual to obtain PCA-derived functional gradients. Notably, the U×S from SVD is equivalent to principal components of eigenvector decomposition.
b. <u>Diffusion Embedding (DE)</u>: This method, a widely used nonlinear dimensionality reduction algorithm, has been used in a recent study demonstrating the major connectome hierarchical systems in both human and non-human primate brains (Margulies et al., 2016). Mathematical details of this algorithm can be found elsewhere (Coifman and Lafon, 2006; Langs et al., 2016, 2014; Margulies et al., 2016; Vos de Wael et al., 2020). Briefly, this algorithm starts with a calculation of an affinity or similarity matrix between given data points. In our case, each data point represents a specific cortical area in the brain, and every cell in the affinity matrix (size: 998×998) refers to the extent of how strong functional connectivity is formed between two brain areas (in case of time series data) or how similar the functional connectivity profiles of two brain areas are (in case of functional connectivity data). For the latter, the functional connectivity matrix can be row-wise thresholded before building an affinity matrix. After rowwise thresholding, the connectivity vector at each brain area becomes sparse, which can affect the similarity calculation. To address this issue, we followed the previous approach (Margulies et al., 2016) to employ cosine similarity as a main distance metric in this study. After calculating the similarity matrix, the DE converted it into a transition probability map (*p*) between data points and estimated its power *p^t^*, which represents the Markov-chain diffuse process evolving as the time *t* along the brain graph (*i.e*., brain nodes linked by functional connectivity). Here, we used 0 (=default setting) for *t*, following the previous study (Margulies et al., 2016; Vos de Wael et al., 2020). By doing so, the algorithm can quantify diffusion distances between cortical areas, which can capture local embedding of a given brain graph based on the eigenvectors and eigenvalues of a diffusion operator.
c. <u>Laplacian Eigenmaps (LE)</u>: This nonlinear dimensionality reduction algorithm has been employed in multiple neuroimaging studies (Haak et al., 2018; Marquand et al., 2017). Similarly with DE, the input to this algorithm was an affinity matrix (with cosine similarity but also eta^2^ similarity, following the previous study, Haak et al., 2018). Minimization of a cost function based on this affinity graph ensures that points close to each other in the original data space are mapped close to each other in the low-dimensional manifold, thereby preserving local distances. LE achieves this goal by calculating the graph Laplacian (L=D[Degree matrix]-A[affinity matrix]) and solving its generalized eigenvalue problem (Lg = λDg where the eigenvectors g_k_ correspond to the *m* smallest eigenvalues λ_k_). Again, the affinity matrix can be constructed directly from time series (998×2380) or based on a thresholded connectivity matrix (998×998).

### 2. Template generation and alignment

As the order of identified gradients and the direction of their signs from dimensionality reduction algorithms are data- and algorithm-specific, the results are often not matched. This mismatch occurs between subjects (even when using the same method), between methods (even when applying to the same subject) and also between test-retest datasets (even if applying the same method to the same subject but with different sessions of data). To make them comparable, a post-hoc matching process to align identified gradients to a reference template is required. To this end, we constructed a group-level gradient template that consists of 250 components (which accounts for 100% of data variability across all algorithms). We first extracted individualized gradient maps (each 250×998), applying PCA to the functional time-series. We then stacked the gradients from all subjects (n=209) into a large 2D matrix (52,250 [=250×209]×998), and performed PCA again on this matrix. This generated a set of group-level gradient templates (250×998), which were used as a reference to which all individual maps were aligned using Procrustes transformation (Wang and Mahadevan, 2008). Of note, *Analyses-2 to −4* used this PCA-derived group-level template, whereas *Analysis-1* relied on the templates directly from each algorithm to evaluate algorithm-specific gradient profiles.

### 3. Between-/within-algorithm reproducibility

One of the important criteria in developing a robust biomarker is reproducibility. To this end, we tested two reproducibility aspects. First, we calculated the within-subject cross-algorithm gradient similarity between PCA, DE and LE. Second, we also calculated within-algorithm gradient similarity between discovery and replication datasets. For each, we examined the reproducibility along the gradient order. To evaluate the effect of a post-hoc gradient matching process, we systematically assessed those reproducibility measures before and after the Procrustes alignment.

### Analysis-2. Reliability evaluation across different parameter setups

We assessed reliability of gradient measures based on both univariate and multivariate statistics, namely intraclass correlation coefficient (ICC, Shrout and Fleiss, 1979) and discriminability (Bridgeford et al., 2020). Briefly, ICC is a statistic defined as the between-subject variability divided by the sum of within- and between-subject variability. While this is one of the widely used reliability metrics, it allows only for univariate items and is valid only under the Gaussian assumption, thus any violation against this condition challenges its interpretation. To fill these gaps, we also included the discriminability (Bridgeford et al., 2020). This recently proposed reliability index is nonparametric, thus requiring no distribution assumption, capable of taking into account for multivariate information and, most importantly, could provide an upper bound on the predictive accuracy of any classification task in unseen data. Statistical definition, theoretical background and validation of this metric can be found in (Bridgeford et al., 2020). Briefly, this measure quantifies the degree to which multiple measurements of the same subject are more similar to one another than they are to other subjects. To do this, it computes the distance between all pairs of subjects, and calculates the fraction of time that a within-subject distance is smaller than between-subject distance. The average of this fraction is referred to the discriminability.

Using these two indices, we have systematically investigated the reliability of functional gradients. Notably, we assessed which combination of dimensionality reduction algorithms and input data types provides the highest reliability, by counting the number of resulting gradients with ICC greater than 0.5. We then visualized the whole brain ICC of gradients from that selected parameter combination and compared them between the algorithms. We also sorted out the ICC spatial patterns based on the established functional community atlas (Yeo et al., 2011) to see which brain network reveals particularly high or low reliability. Moreover, to decompose the sources of ICC, we separately calculated within- and between-subject Euclidean distances in the gradient space and see which one explains more dominantly those high ICC values. Lastly, the discriminability index was assessed across the algorithms, stratified based on the input data types (*i.e*., time-series, differently thresholded functional connectivity matrices).

### Analysis-3. Reliability evaluation across different R-fMRI time-series length

Apart from the algorithm used and input data type, another critical factor affecting the quality of extracted gradients is an amount of the data available (*i.e*., a time-series length of R-fMRI). Given that most clinical studies rely on relatively limited amount of R-fMRI data due to practical challenges, figuring out a lower bound of the scan time to obtain reasonable reliability and sensitivity in detecting behavioral association will have a direct impact on prospective data collection. To address this question, we benefited from the densely sampled individual data from MSC, where 9 subjects are available, each having 10 different sessions of R-fMRI. The targeted time-series lengths were 5-, 10-, 20-, 30-, 40- and 50mins. To generate these fMRI time-series lengths, we randomly chose a contiguous segment(s) of R-fMRI time-series as the same length as a targeted time length among the entire 300mins data and merged them if needed. For instance, to make 40mins data, we selected one 30mins contiguous segment keeping the original volume order (please recall that one session data of MSC is 30min) and another 10mins of data and merged them. This strategy to choose ‘contiguous’ volume segments was to keep any inherent properties of the original data related to its stationarity, while randomly aggregating the data. This process was performed two times separately, in order to make test-retest datasets. Once we generated these different lengths of fMRI data, we systematically applied the three dimensionality reduction algorithms to evaluate their reliability as a function of an amount of the data.

In this analysis, we focused on the 2^nd^ PCA-derived gradient, given its highest ICC among multiple combinations of parameters and different gradient order. We iterated the above random data generation process and reliability calculation 10 times and reported the averaged results. We also categorized the resulting ICC values according to the widely accepted interpretation guideline (ICC≤0.4: fair, 0.4<ICC≤0.6: moderate, 0.6<ICC≤0.8: substantial, 0.8<ICC<1: almost perfect; Landis and Koch, 1977) as well as in terms of canonical functional communities to display the network-stratified ICC patterns across different algorithms. Finally, we performed the same analysis assessing the effect of amount of the data based on discriminability.

### Analysis-4. Prediction framework based on canonical correlation

Because of their statistical definition, both ICC and discriminability serve as indicators for a degree of how unique individual information the given measure retains to distinguish it from the group of items or subjects (Zuo and Xing, 2014). Hence, they are not only reliability metrics but also can be used as a marker for inferring the upper bound of prediction power of a given metric. For instance, if the reliability is low, there is less probability for this measure to predict independent phenotypic data, since the measure (here, a gradient) per se is already less individually distinguishable. To explicitly test this relationship (*i.e*., reliability *vs*. prediction accuracy) for gradients, we performed a gradient-based prediction analysis for phenotypic scores provided in HCP and related the prediction accuracy to reliability of the gradients.

### 1. Profiling of HCP phenotypic scores based on their covariance matrix and factor analysis

The 65 HCP phenotypic scores targeted in this study are categorized into 6 different domains including alertness, cognition, emotion, personality, sensory and psychiatric/life functions (Barch et al., 2013). Given their high interdependency (see **Supplementary Figure 1A**), it is tempting to hypothesize the existence of underlying latent factors. To test this hypothesis, we first applied an exploratory factor analysis on the 209 (subjects) × 65 (phenotypic scores) matrix and obtained factor scores to see overall individual phenotypic patterns. We then used these factor scores as a responder in following prediction analyses.

### 2. *Prediction framework* (Supplementary Figure 2)

The predictor was a single gradient map derived from a specific dimensionality reduction algorithm and input data type (*e.g*., PCA on a 95% thresholded connectivity matrix) using 30mins of R-fMRI data across individuals (size=209×998 [subject by brain areas]). The responder was a single phenotypic score of the same individuals (size=209×1; either factor score or raw phenotypic score). While previous analyses targeted 250 gradients in total, here we focused on only the first 100 gradients because the later gradients explained less than 1% of the variance of the original connectome data. We used a canonical correlation analysis (CCA) to associate the two sets of variables (*i.e*., predictors and responder, McIntosh and Mišić, 2013; Smith et al., 2015; Wang et al., 2018). Briefly, CCA – a generalized multivariate correlation approach – finds linear combinations of the variables in each of two multivariate sets such that the two sets make the best correlation with each other. As a result, CCA provides canonical coefficients (weights for linear combinations) which, in the prediction context, become trainable parameters that will be applied to the unseen test cases. Before prediction, we first performed PCA on the gradient matrix (predictor; 209×998 [subject by brain areas]) to reduce its original high dimensionality into 209×X (X = # of the components that can explain >90% of variability of an original matrix). This dimensionality-reduced gradient matrix and the phenotypic score were then fed into CCA to find canonical coefficients. After learning both PCA and CCA coefficients of gradients from the training data (discovery), we applied the PCA coefficients to raw gradient scores of the test cases (test-1), and then the CCA coefficients to these reduced features, and finally performed an inverse mapping of PCA in order to reconstruct phenotypic scores. The prediction accuracy was measured using Spearman correlation between the predicted phenotypic scores and the original scores.

We created the above prediction framework 100 [# of targeted gradients] × 3 [# of factor] or 65 [# of raw phenotypic scores] times. This group of predictions was again iteratively performed across all pairs of 3 different algorithms × 11 input data types (time series and differently thresholded functional connectivity matrices). Increased Type-I errors due to multiple prediction accuracy calculations (*i.e*., Spearman correlation) were controlled by false discovery rate (FDR) at 5% (Benjamini and Hochberg, 1995). We counted the number of gradients showing FDR-survived significant prediction across different input data types at each algorithm (*i.e*., PCA, DE and LE) to assess the parameter combination providing the most predictive gradient markers. All prediction analyses were repeated using a second independent dataset (test-2) for reproducibility of findings.

### 3. Relationship between prediction accuracy and reliability

Once we obtained a prediction accuracy table (100[# of targeted gradients] × 3 or 65[# of factor or phenotypic scores]), we averaged the accuracy values at each row (=each gradient) to measure a general prediction power across phenotypic scores. We then correlated this averaged prediction accuracy and reliability (whole-brain averaged ICC and discriminability) based on Spearman correlation. We tested this prediction-reliability correlation across all combinations of the algorithms and input data types.

### 4. The effect of R-fMRI time-series length on prediction

We evaluated the effect of R-fMRI time-series length on prediction accuracy. As done in *Analysis-3*, we selected the gradients from PCA applied to the 95% thresholded functional connectivity, given its highest reliability. Because the MSC data has only 9 subjects, it was not enough to find generalizable CCA coefficients for prediction. Thus, we instead used the HCP dataset, randomly choosing the segments of R-fMRI volumes among one-hour data, by varying the time-series length. We created different lengths (*i.e*., 5-, 10-, 20-, and 30mins) of test-retest time-series data from both training and test-1 cases, and conducted the same reliability/prediction analyses as done in *Analysis-3 and −4*. We also performed a prediction analysis using 50mins data to see if the accuracy keeps improving without making a saturation.

### 5. Comparison with conventional edge-based connectome prediction

Finally, to assess the unique strength of low-dimensional gradient approaches compared to the conventional methods, we selected the recently proposed, connectome-based predictive modeling (Shen et al., 2017, YaleMRRC/CPM) as a reference to compare. This simple yet powerful method directly takes a connectivity matrix of individuals as an input, trains a robust regression after selecting only predictive edges (connectivity) and tests unseen samples in a cross-validation setting. The only parameters that can be tuned in CPM is the alpha threshold for the selection of edges showing a high correlation with a given responder. To make a fair comparison, therefore, we systematically varied the threshold of CPM between 0.001 and 0.05 with every 0.0025 interval and aggregated only the significant predictions (after FDR corrections). We tested the CPM based on factor scores, and compared the results to those of gradients.

## Results

### Gradient extraction, alignment, and reproducibility tests

Comparison of the connectivity gradients generated using PCA, DE, and LE (**Figure 1**) suggested that similarities exist among the algorithms, though primarily for those explaining the highest amounts of variance. Specifically, the first four gradients generated by the three algorithms were highly similar (*e.g*., spatial correlation across the 4 gradients: 0.86±0.11 [PCA-DE], 0.97±0.02 [DE-LE], 0.90±0.07 [PCA-LE]), and the later components exhibited a rapid drop-off in similarity across these methods. Given the expectation that the gradient components explaining less variance would be more sensitive to the choice of algorithm, we matched components across the three algorithms using Procrustes alignment (Wang and Mahadevan, 2008). This procedure suggested a notably higher degree of similarity in the results (at least for 10 components as shown in **Figure 1C**). Importantly, we found that those gradient components having a high cross-algorithm similarity also exhibited a high degree of reproducibility across the samples (replication-1) as well.

**Figure 1.**
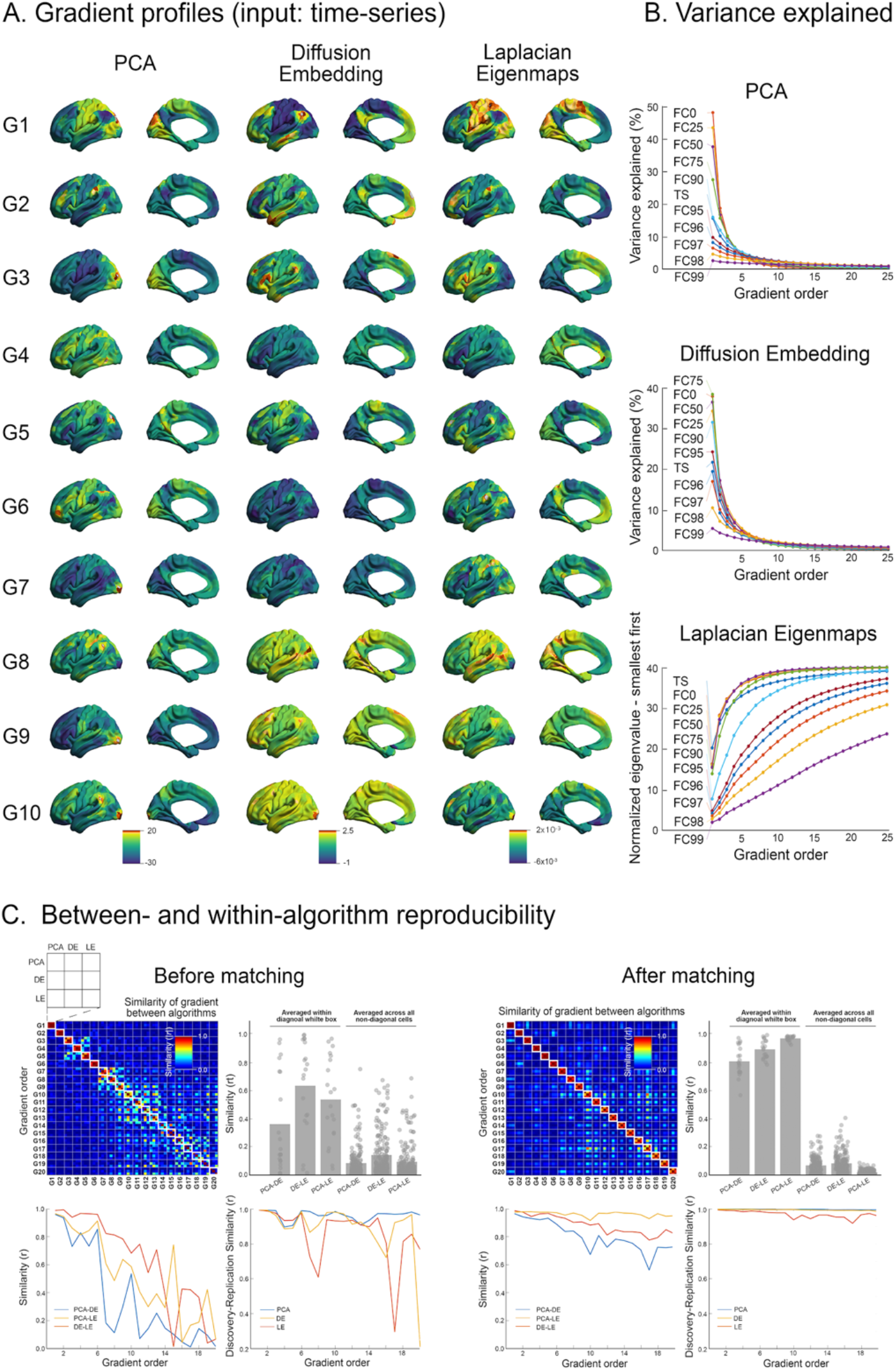
Profiling of functional gradient and its reproducibility. **A)** Mapping of the first 10 gradients directly from time series data (PCA, DE, and LE in order) before alignment. **B)** Relative proportion of variance explained for PCA and DE as a function of gradient order (x-axis) and threshold (color-coded). Please note that LE has algorithmically different principles in terms of ordering the components (selecting the smallest eigenvalues first), thus showing upside-down flipped curves compared to those of PCA and DE. **C)** The summary of cross algorithm similarity of the first 20 gradient maps was shown before (left) and after (right) matching with Procrustes transformation. In the right bottom corner, the test-retest reproducibility of the gradients is also present between different samples before and after alignment. The first few gradients (*e.g*., 1-6 gradients) are similar to one another, regardless of whether matching was used, and that for the later gradients, a gradient alignment dramatically improved both the cross-algorithm and cross-sample reproducibility.

### Reliability evaluation across different parameter setups

Regardless of the algorithm used, ICC was highest for those that explain a greater proportion of the variance (*i.e*., the lower order gradients). This is consistent with our findings that lower order gradients are more stable and replicable within a subject – properties that would be expected to yield higher test-retest reliability. Second, generally we found an advantage for using threshold matrices over time-series data, and particularly more conservative thresholds (*e.g*., >90%). Of note, as one would expect, too conservative thresholding (>98%) turned out to actually reduce reliability, suggesting that excessive thresholding can remove important individual variations in connectome.

In **Figure 2**, to illustrate key points, we depicted the reliability maps derived from 95% threshold functional connectivity. Yet, the findings were generalizable to other combinations of parameters, which are presented in **Supplementary Figure 3**. First, when the vertices are sorted into networks, we found that those in the dorsal attention, frontoparietal and default mode system have higher ICC than the rest of the brain (two sample t-tests between these two network systems: p<0.001, t=17.2 for PCA; p<0.001, t=14.9 for DE; p<0.001, t=19.7) – regardless of the algorithm employed (PCA, DE, LE). Examination of contributing sources of variation to ICC revealed that these high ICCs are primarily derived from higher between-subject variability of those gradients, rather than lower within-subject variability. Overall, discriminability was highest for the PCA, and lowest for LE. For the gradients from LE, given that previous studies employed the eta^2^ similarity (Haak et al., 2018; Marquand et al., 2017) for the affinity matrix calculation, we also evaluated the reliability based on this approach, and found slightly decreased reliability compared to the ones from the cosine similarity (**Supplementary Figure 4**).

**Figure 2.**
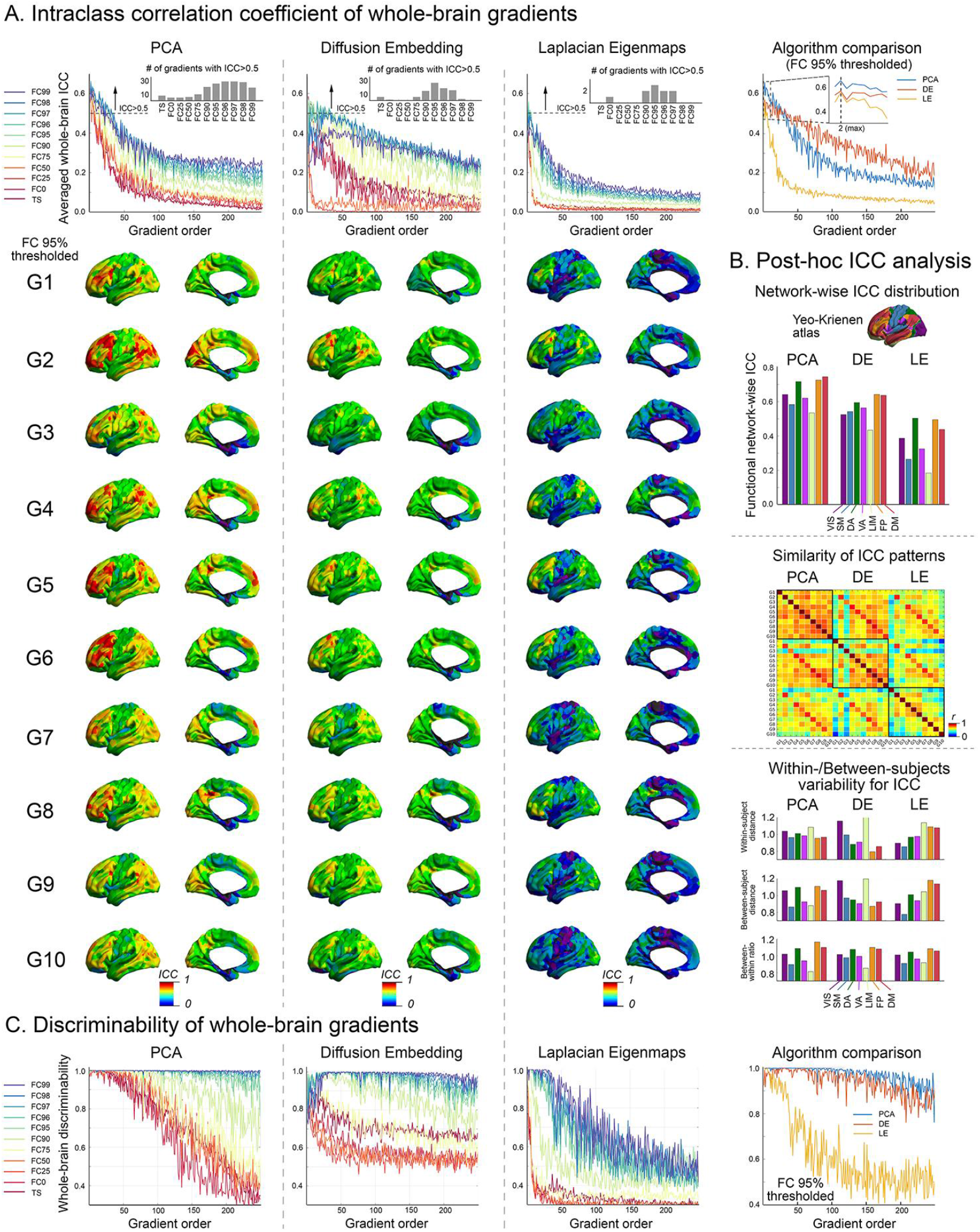
Reliability of functional gradients. **A)** *Upper*: The whole-brain mean intraclass correlation coefficient (ICC) is plotted across the thresholding approach (color-coded) and as a function of the gradient order (x-axis) for the three dimensionality reduction algorithms. The inset at each graph presents the number of gradients with >0.5 ICC (all algorithms commonly show the peaks in the range of 95-97%). The rightmost graph shows cross-algorithm comparisons of ICC at 95% thresholding. *Bottom*: The spatial distribution of the ICC for the first 10 matched gradient components is mapped on the whole-brain cortex. **B)** *Top*: At each algorithm, averaged ICC across 1-10 gradients was sorted out based on a canonical functional network atlas (Yeo et al., 2011). *Middle*: The similarity of whole-brain ICC patterns across the gradients is shown. Note that the diagonal components represent within-algorithm similarity, whereas the off-diagonal shows between-algorithm similarity. *Bottom*: The two sources contributing to ICC, namely within-subject and between-subject variability, are separately computed across the three algorithms. Note that high ICC is generally driven by high between-subject variability rather than low within-subject variability. **C)** The discriminability of gradients across the algorithms as a function of gradient order (x-axis) and thresholding approach used (color-coded).

### Reliability evaluation across different time-series lengths

Here, we examined the relationship between R-fMRI time-series length and gradient reliability, both at the vertex and whole-cortex levels. To accomplish this goal, we leveraged the Midnight Scan Club data (Gordon et al., 2017), which allowed to create two subsets of data of varying amount of R-fMRI data per participants (*i.e*., 5-, 10-, 20-, 30-, 40-, 50 minutes). Consistent with prior findings (Birn et al., 2013; Noble et al., 2017), we found that regardless of the connectivity gradient algorithm or thresholding strategy employed, vertex-wise reliability (indexed by ICC) progressively increased with the amount of data available. For the purposes of clarity, we focused our reporting on the findings from the PCA/95% thresholding strategy (**Figure 3**), which were superior in terms of reliability. We found that at 20 minutes, >50% of vertices achieved at least moderate reliability (ICC>0.4) and >30% of the vertices reached the classification of substantial (ICC>0.6). Of note, the number of vertices achieving reliability that would be classified as “substantial” or “almost perfect” more than doubled when increasing from 5 to 20 minutes, and from 20 to 50 minutes. Importantly, when we sorted vertices by network, we found that those in the default, attention and frontoparietal networks tended to be substantially higher than those in the visual, somatomotor and limbic networks across all time-series lengths (two sample t-tests at each length [showing the least significant statistics among 5 different lengths]: p<0.001, t=11.1 for PCA; p<0.02, t=2.43 for DE; p<0.06, t=1.91 for LE).

**Figure 3.**
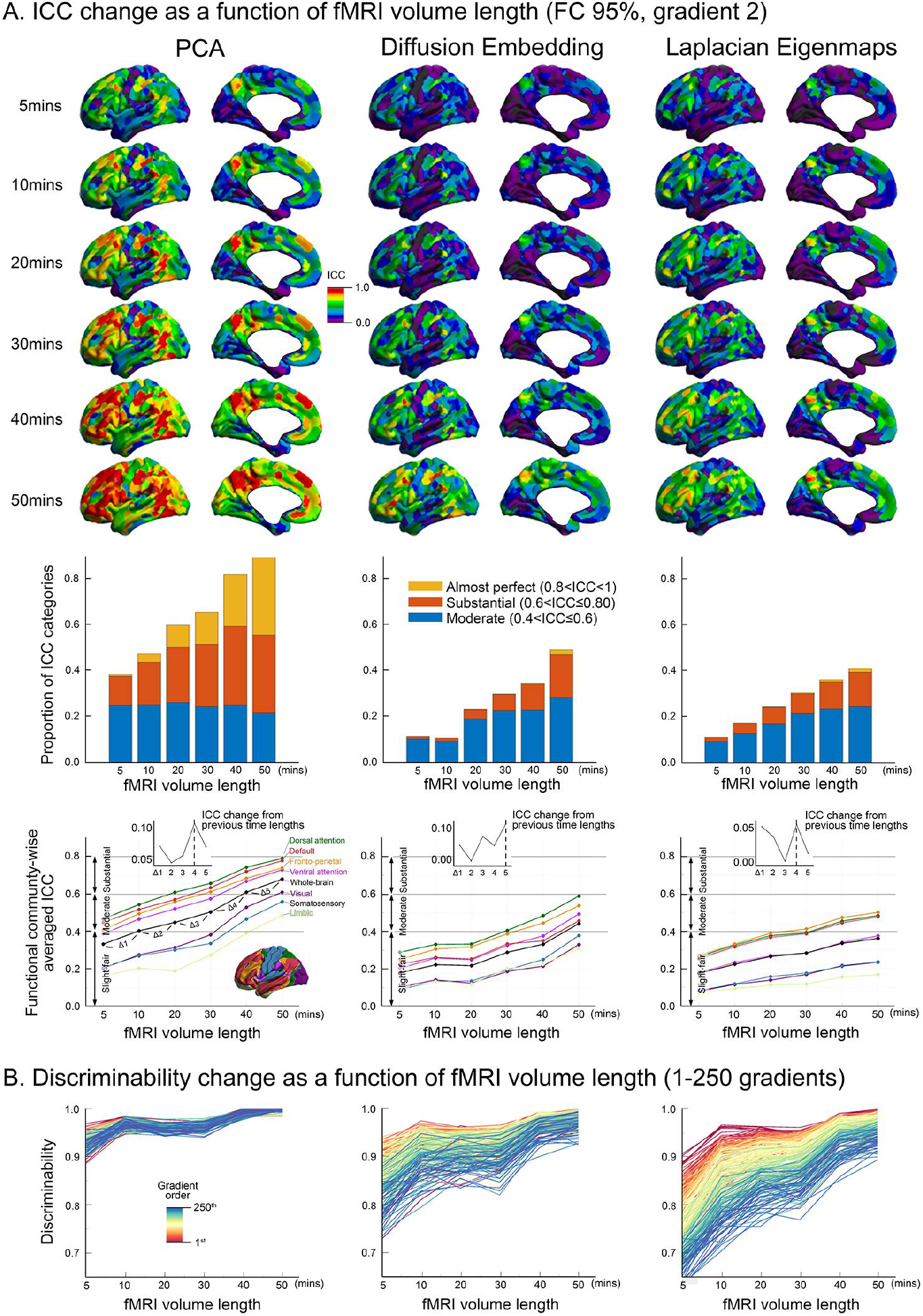
Effects of scan duration on reliability. **A)** *Top*: The changes of ICC across the whole brain are shown as a function of increasing R-fMRI time-series length for the second gradient from the 95% thresholded functional connectivity matrix. Overall, PCA shows the highest regional ICC as increasing time-series length. PCA with only 5 minutes of data produces ICC values comparable to DE with as much as 40 minutes of data. *Middle*: The changes of ICC as a function of time-series length are categorized into widely used ICC interpretation criteria (ICC≤0.4: fair, 0.4<ICC≤0.6: moderate, 0.6<ICC≤0.8: substantial, 0.8<ICC<1: almost perfect; (Landis and Koch, 1977). Here only from the moderate range is shown to focus on relatively acceptable ICC values. Please note that from 20mins data already >60% of vertices over the whole brain present ICC>0.4. DE and LE follow PCA in order. *Bottom*: As done in *Figure 2*, the data amount-dependent whole-brain ICC changes were stratified into 7 functional communities (Yeo et al., 2011) and shown across the algorithms. To assess how the ICC changes as every 10mins-length increase, we computed the difference of ICC between the current time length and the previous one. The peak of this ICC changes occurs normally in longer time-series (*e.g*., 40-50 mins). **B)** The discriminability as a function of increasing timeseries length is present across the full gradients. Note that in 20mins or more data, all PCA gradients are highly discriminable, whereas there exist a range of discriminability values across gradients for DE and LE.

Our examination of discriminability suggested a more complex relationship across different orders of gradients. In all algorithms, the gradients explaining larger data variance tended to exhibit greater discriminability than those ones explaining smaller variance, with differences in discriminability among gradients decreasing with the amount of data. Of note, regardless of the algorithm employed, increases in discriminability as a function of fMRI time-series length were prominent when increasing from 5- to 10-minutes, and from 30- to 40-minutes; findings were otherwise relatively steady across durations.

### Gradient-based prediction analysis

We systematically evaluated the prediction ability of functional gradients across different combinations of the algorithms and time-series/connectivity thresholding strategies.

1. *Factor score prediction* (**Supplementary Figure 1**): Before a prediction analysis, we first profiled the patterns of 65 HCP phenotypic scores. In exploratory factor analysis, we focused on the 3 factor-model because *k*=3 made an elbow point in the graph of variance explained. The resulting three factors were summarized into *i*) <u>externalizing</u> (psychiatric/life function [thought problem, attention problem, rule break, other problems, total problem, ADHD, inattention problem, hyperactivity, antisocial]), *ii*) <u>internalizing</u> (emotion [anger, fear, sadness], social relationship [loneliness, hostility, rejection, perceived stress], personality [neuroticism], psychiatric/life function [withdraw, internalizing, depress, anxiety, avoid]), and *iii*) <u>general cognitive function</u> (cognition [fluid intelligence, language reading/comprehension, spatial orientation processing, verbal episodic memory], emotion [emotional recognition]). From these factors, we extracted individual scores to enter into a prediction framework as responders.

In the test-1 dataset, when using gradients from PCA and 95% thresholded connectivity matrix, the factor-3 representing global cognitive functions was generally well predicted across many gradients (# of gradients showing significance=61 out of 100 gradients after FDR correction), while the other twos (*i.e*., externalizing, internalizing) showed significance in only a few gradients (**Figure 4A**). This pattern was largely replicated across other combinations of algorithms and input data types as well (**Figure 5A**), suggesting a strong association of functional gradients towards general cognitive performances. Notably, when associating this prediction accuracy to reliability (*i.e*., ICC and discriminability), it showed strong positive correlations (**Figure 4A**), reflecting a clear advantage of assessing reliability in inferring a phenotypic prediction power. This prediction-reliability relationship was consistently found in the second independent dataset (test-2) as well.

**Figure 4.**
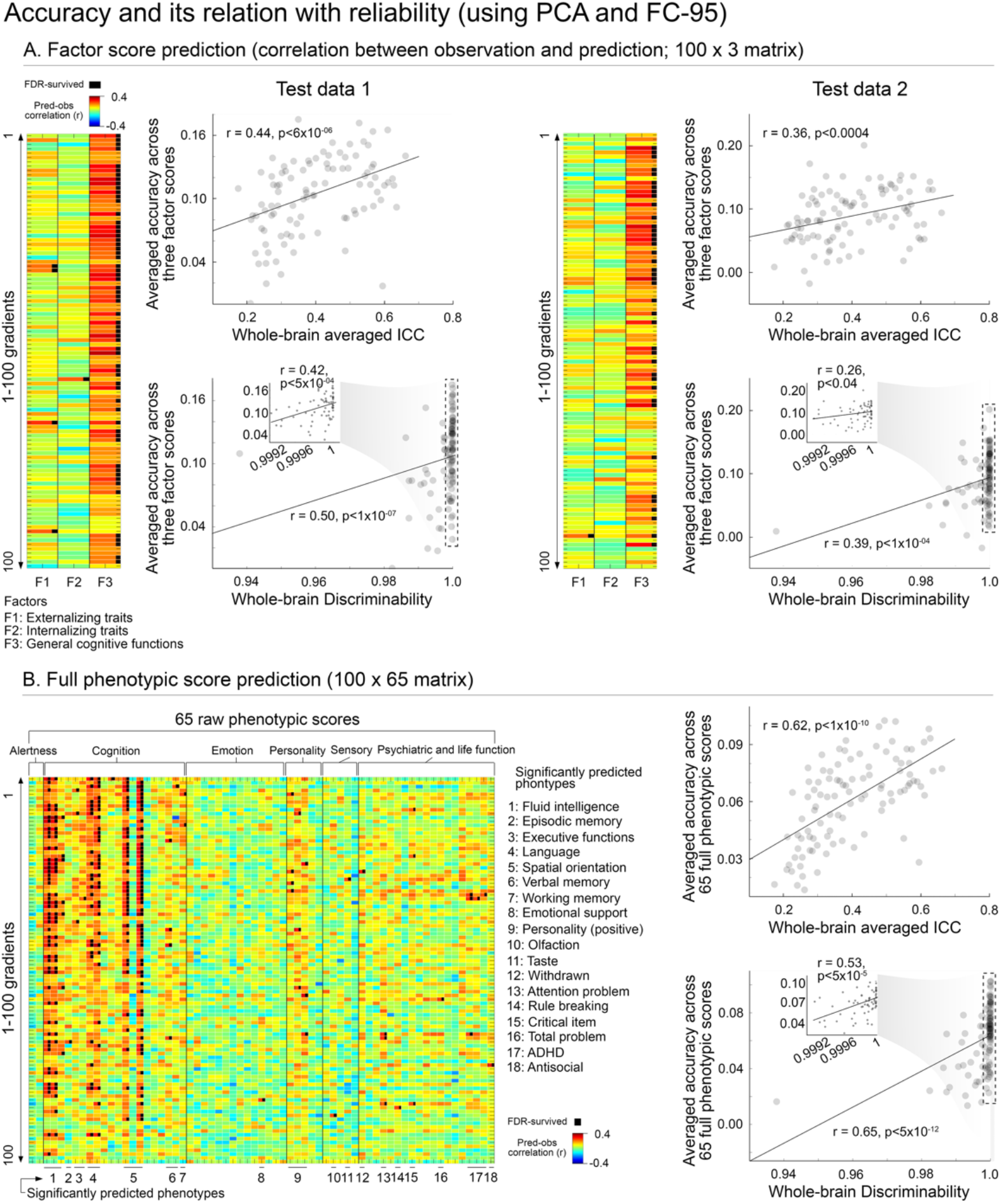
CCA-based behavioral prediction and its relationship to reliability. **A)** The results of three-factor score prediction using gradients from PCA and 95% thresholded functional connectivity are present (left: test-1 and right: test-2). Each cell in the left matrix represents the prediction accuracy (Spearman correlation between observed and predicted scores) of 100(# of predictors [gradients]) × 3(# of factors [externalizing, internalizing and general cognitive functions) tests. The black dots in the matrix indicate FDR-survived predictions. Please note that the most FDR-significant predictions occur from general cognitive functions, and only few in the externalizing and internalizing symptoms. The next scatter plots present the relationship between reliability (top: ICC, bottom: discriminability) and prediction accuracy (*i.e*., row-wise averaged *r* values from the left matrix). Both show a strong positive relationship, suggesting that more reliable the gradient is, higher prediction power it has for unseen phenotypic scores. Notably, in the discriminability plot (bottom), most dots are positioned in the rightmost side. However, if the rightmost side (the range for nearly perfect discriminability) is stretched out, the discriminability is still correlated to the prediction accuracy as shown in the inset. Virtually identical results were replicated in the test-2 datasets. **B)** The same format of the results but for the entire 65 phenotypic scores (belonging to the 6 different domains) were presented. Again, the black dots indicate the FDR-significant prediction, and the predicted 18 different phenotypic scores were listed next to the matrix. The rightmost scatter plots are the relationships for reliability (top: ICC, bottom: discriminability) and averaged prediction accuracy.

**Figure 5.**
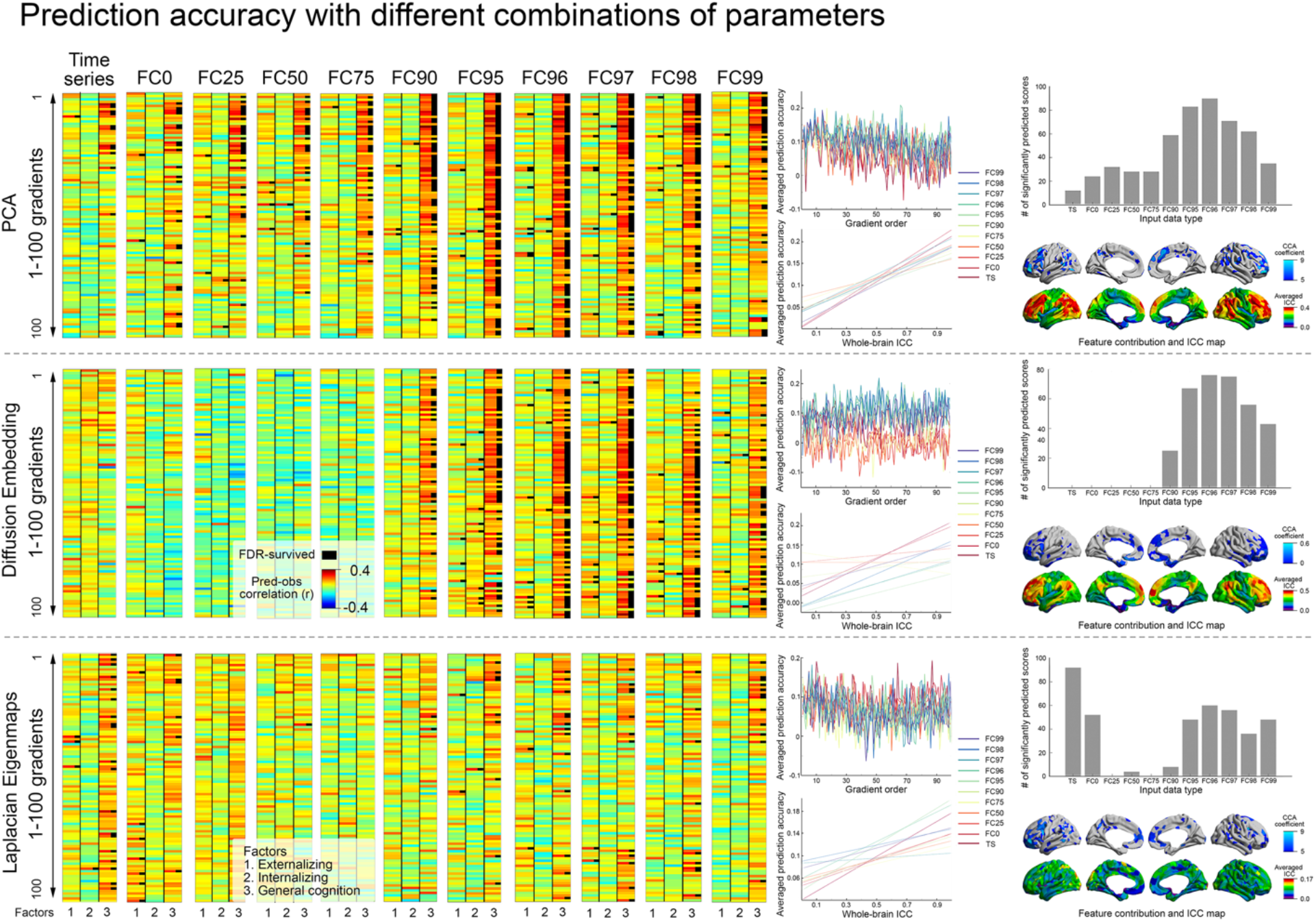
Comparison of behavioral prediction across algorithms and input data representation. *Left*: The group of matrices (size: 100×3×11 [# of gradients×# of factors×# of input data types]) presents the distribution of three-factors prediction accuracy as a function of input data types (*i.e*., time series, 0, 25, 50, 75, 90, 95, 96, 97, 98, 99% thresholded from left to right) and the gradient algorithm used (PCA, DE, LE). Note that for both PCA and DE, the predictive accuracy is maximized for the upper thresholds (95%-97%). *Middle*: Averaged prediction accuracy is presented across different input data types (color-coded) as a function of gradient order (top) and as a function of whole-brain ICC (bottom). As the gradient order gets higher, prediction accuracy gradually decreases. Notably this pattern was distinct across the input data types, with a tendency of less-conservatively thresholded functional connectivity (*e.g*., 0-50%) showing generally lower prediction accuracy compared to the highly thresholded connectivity input. As whole-brain ICC increases, the prediction accuracy accordingly increases. *Right*: In both PCA and DE, the 95-96% thresholding range shows the highest number of FDR-significant prediction gradients, whereas in LE, the time-series based gradients show the highest number of significant predictions. Notably, when mapping the feature contribution from CCA training, its spatial distribution appears highly resembling patterns with the ICC wholebrain profiles regardless of the algorithms used.

Beyond prediction based on a single gradient map, we also tested whether combining multiple gradients can further improve the prediction accuracy. The first step in this process was to identify the most promising subset of gradients to combine, in an unbiased manner. To accomplish this, we split the training cases (*i.e*., discovery dataset) into 5 folds, and performed a within-sample prediction analysis, which provided a set of gradients showing significant factor-score prediction. We combined these gradients into a single, large feature matrix (209× [998× # of gradients]) to enter it to the CCA framework. We then followed the same prediction procedures as above (see ‘*2. Prediction framework*’ in the **Method**) to test unseen cases (replication-1). It is important to note that the test cases were completely isolated from this gradient selection process. The result showed that the combined gradient approach greatly improved prediction accuracy in general cognitive domains from 0.33 (maximum Spearman correlation in the single-gradient based prediction) to 0.41, suggesting the collective power of useful gradients with high predictive ability (see **Figure 6**). On the other hand, the externalizing symptoms showed a decreased performance (from 0.2 to 0.12 Spearman coefficient).

**Figure 6.**
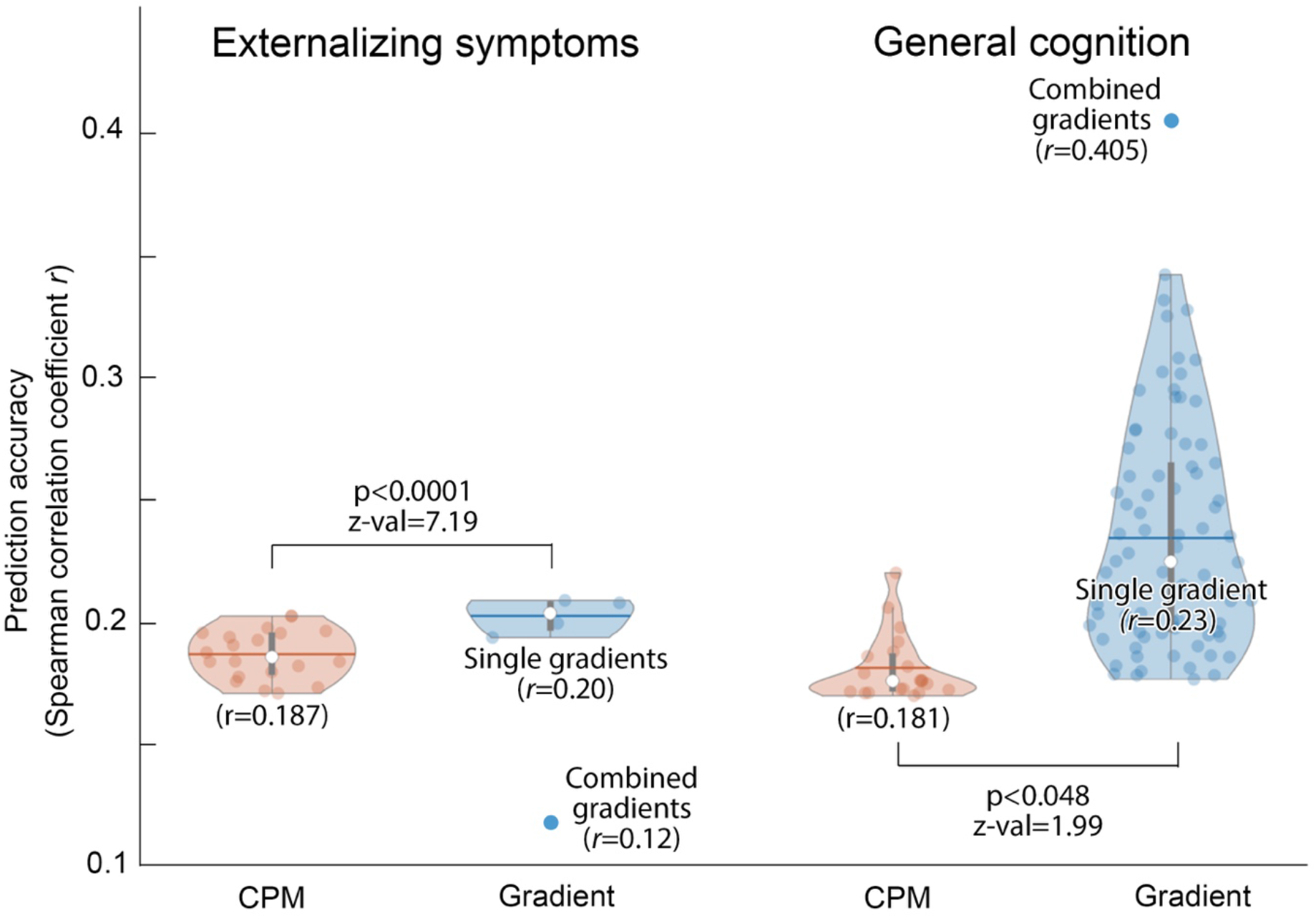
Comparisons of prediction accuracy between functional gradients and connectome-based predictive modeling (CPM) approaches. Only the factor scores showing significance prediction in both gradient and CPM approaches are shown (*i.e*., externalizing symptoms and general cognition domains). Individual pale blue dots in the violin plot represent the significant accuracy among 100 gradients, whereas the orange dots are the accuracy derived from the tests of systematic p-value thresholding for feature selection in CPM. Additionally, an individual isolated blue dot (filled) above or below each gradient result represents the prediction accuracy when using multiple gradients. See ‘*Gradient-based prediction analysis*’ in **Method** and ‘*Fullphenotypic score prediction*’ in **Results** for details. Most accuracy values in the single gradient approach outperform those of CPM, suggesting added values of lowdimensional connectivity representation compared to the pure edge-based analysis. The statistics are based on nonparametric Wilcoxon Rank sum tests comparing the accuracy between the two methods. Combining multiple gradients showed a prediction improvement in the general cognitive domain. The reason why the same multiple-gradient approach yielded a worse accuracy in the externalizing symptoms may be likely because of less generalizability about significantly predictive gradients for externalizing symptoms between training and test datasets.

In part, this may suggest that our strategy to select gradients for combination was suboptimal; toward this point, had we selected the individual gradients that performed best in the test dataset, and combined them, prediction would have gone up to 0.32 for externalizing. Alternatively, it may reflect the fact the effects for the psychiatric factors were notably smaller than those observed for the cognitive −possibly due to the fact that the HCP samples are largely neurotypical subjects. Related to this point, recent work has highlighted the suboptimal nature of psychiatric tools such as the Adult Self Report in non-psychiatric samples (Alexander et al., 2020).

Finally, we assessed the difference between ICC (univariate) and discriminability (multivariate) in terms of their ability to infer a phenotypic prediction. To this end, we constructed a precision-recall curve at each reliability measure, by which we could compare how much their reliability can screen only ‘FDR-survived’ significant phenotypic predictions. Again, here we used the prediction result from PCA applied on 95% thresholded functional connectivity matrix. This analysis demonstrated no statistical differences between the two measures (**Supplementary Figure 5**), although ICC revealed the FDR-survived significant prediction even in the relatively lower, arbitrary thresholds (ICC=0.2-0.3), whereas in discriminability only nearly perfect thresholds (=1) suggested FDR significances, which may serve as a practically more useful criterion, given its non-arbitrariness.

*2. Full phenotypic score prediction*: Similar prediction results were observed in raw 65 phenotypic scores as well (**Figure 4B;** note that the result was based on PCA applied on the 95% thresholded functional connectivity). Indeed, most gradients showing FDR significance were found in the categories of cognition and psychiatric/life function, and much less in other domains. Specifically, those scores showing at least one significant prediction were found: in the *cognition* domain, fluid intelligence, episodic memory, executive functions, language, spatial orientation, verbal/working memory; in *emotion*, emotional support; in *personality*, positive traits (agreeableness, openness); in *sensory*, olfaction and taste; in *psychiatric and life function*, withdrawn, attention problem, rule breaking, critical item, total problem, ADHD and antisocial. As in the factor-score based analysis, the averaged prediction accuracy across these phenotypes was positively correlated to ICC and discriminability, emphasizing utility of reliability.

When expanding this analysis towards other combinations of different algorithms and time-series/connectivity thresholding strategies (**Figure 5**), PCA-based gradients showed overall higher prediction rates compared to other methods. Moreover, the 95-97% of thresholding appears to be the most predictive range in both PCA and DE, as similarly found in their reliability profiles. Notably, when mapping the canonical coefficients learned from training across all gradients showing significant prediction, the spatial patterns across the whole brain highly resembled those areas showing high vertex-wise ICC, suggesting a strong relationship between reliability and individual prediction in the local brain areas.

*3. The effect of time-series length on prediction*: Similar to what has been done in *Analysis-3*, we systematically varied the time-series length to construct functional gradients and investigated the corresponding changes of prediction accuracy. Exemplifying the results based on PCA applied on 95% thresholded functional connectivity (**Figure 7**), we demonstrated significant effects of time-series length. Specifically, while the increase of the length made a monotonous increase of prediction performances, particularly 10-to-20mins data length change yielded the largest increase of a prediction performance, whereas it rather remains stable during 5-10mins and 20-30mins time increase.

**Figure 7.**
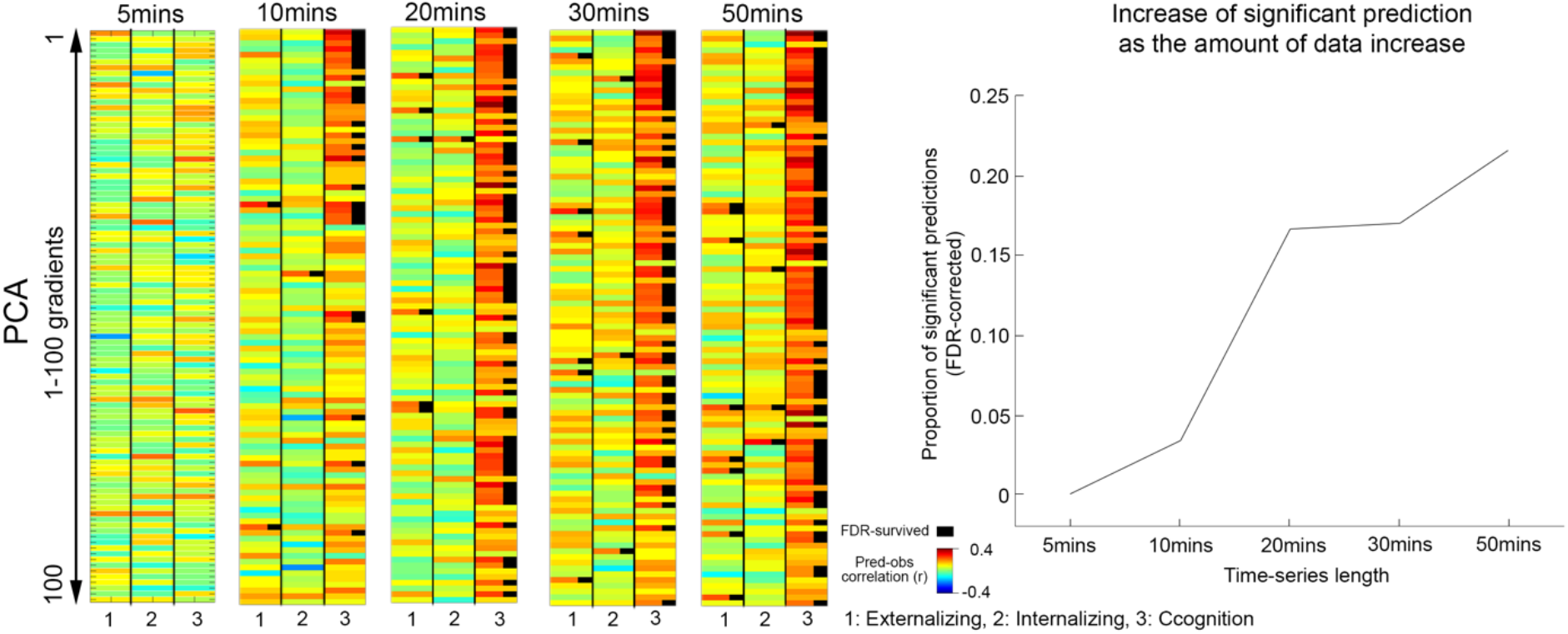
The effect of the data amount (*i.e*., time-series length) on phenotypic outcome prediction and its relationship to reliability. *Left*: The prediction accuracy for factor scores based on gradients from PCA and 95% thresholded functional connectivity is shown as a function of different time-series length (5, 10, 20, 30, 50mins). *Right*: As the amount of the data gets larger, the number of FDR-significant prediction cases also monotonously increases.

*4. Comparison with conventional edge-based prediction*: The CPM analysis consistently yielded significant phenotypic predictions across almost all alpha thresholds in externalizing symptoms and general cognition domains (Spearman coefficients ranged from *r*=0.17 [p<0.013] to r=0.20 [p<0.003] for externalizing; and *r*=0.17 [p<0.014] to r=0.22 [p<0.0014] for cognition; all p values survived the FDR correction), though none for internalizing. Yet, the accuracy level was significantly lower compared to the gradient approaches (see **Figure 6**) in both phenotypic scores, as indicated by Wilcoxon rank sum tests between CPM and gradients (p<0.001 for externalizing and p<0.048 for cognition), suggesting added values of low-dimensional functional connectivity representation.

## Discussion

The present work evaluated the suitability of connectivity gradients for cognitive and psychiatric biomarker discovery. We identified a benchmark set of parameter selections that maximizes reliability and as a result, enhance the ability to predict features of a specific individual. Our analyses focused on three key factors, (*i*) reproducibility, (*ii*) reliability and (*iii*) predictive power, and explored how they change depending on the threshold and type of functional similarity data, how gradients are extracted, and the amount of data. While there are many factors that can potentially determine reliability, we found that certain sets of the analytical strategies for the calculation of gradients were more useful in the context of biomarker discovery. These include *i*) using a linear dimensionality reduction algorithm (*e.g*. PCA), *ii*) utilizing the gradients that explain a greater amount of the variance of the original data, *iii*) extracting gradients using more conservatively thresholded functional connectivity matrices and *iv*) focusing on more reliable and predictively powerful high-order transmodal systems rather than low-level primary sensory systems. Notably, while our findings experimentally support these recommendations, future work should tailor them, depending on the analysis goal of a given study.

Our study examined two different aspects of reproducibility, one focused on cross-algorithm similarity and the other on the replication of findings in different sample data (replication-1). A well-established caution that exists for many dimensionality reduction techniques, is that component orders and directionality (either positive or negative) can be sensitive to minute differences in data. As a result, testing reproducibility directly on the raw functional gradients across different methods and thresholding strategies may not be appropriate. Examination of the raw functional gradients across algorithms suggested significant variation, particularly in the gradients accounting for less variance. Importantly, our findings demonstrated the utility of Procrustes alignment in matching gradients across techniques, parameters and subjects to enable meaningful comparisons, and demonstrated a high degree of cross-algorithm agreement once properly aligned. Another noteworthy point is that the algorithms differed in the amount of variance explained along the gradient order. By the algorithmic nature of PCA, its lowest order gradients (*e.g*., the first few primary components) explain the highest amount of variance, while the nonlinear algorithms exhibited variances that are more evenly explained by the gradients along the order (especially when the gradients were based on more conservatively thresholded functional connectivity [98-99%]). This may explain why measures of reliability tended to drop off more rapidly as gradient order increased with PCA than with its nonlinear alternatives (**Figure 2A**). It also emphasizes the value of thresholding in increasing reproducibility across algorithms.

It is important to note that although our findings favor PCA with respect to reliability and predictive validity of individual gradients, this does not negate prior findings with the more sophisticated, nonlinear algorithms examined (*i.e*., diffusion embedding, Laplacian eigenmaps). Our results, however, do suggest the merits of carefully establishing the need for such algorithms over more basic methods (*e.g*., PCA) before their adoption. This consideration may be particularly relevant when gradient techniques are applied to regional brain areas. Indeed, a recent study (Haak et al., 2018) using Laplacian Eigenmaps (LE) found a unique ability of non-linear manifold learning to recapitulate the functional connectivity organizations (*i.e*. connectopies) of visual and somatosensory areas, which was not able to be detected using PCA (applied to un-thresholded data, which we found to be suboptimal in our whole-brain application). Along the same lines, we also found that those nonlinear methods explain greater variance in higher order gradients compared to PCA, suggesting that these approaches have higher sensitivity in probing local, more nonlinearly embedded data structures in the manifold space. The current moment is especially useful to test these methods and better understand costs and benefits given their growing availability in relatively turnkey packages, such as BrainSpace (Vos de Wael et al., 2020). At a minimum, studies using nonlinear algorithms could benefit from including a comparison against linear methods to justify their use in a specific context. As the present work was comprehensive, though not exhaustive, it is undoubtedly possible that variations of these nonlinear algorithms and their parameters may be able to bring performance to be notably in excess of that seen with PCA. Future work would benefit from such demonstrations.

Beyond algorithm selection, we also found that the application of thresholds to correlation matrices and the type of affinity matrix calculation (*e.g*., cosine vs. eta-squared similarity) in gradient analyses play an important role to determine its reliability. While many studies include supplementary analyses that vary the threshold applied to demonstrate the robustness and limitations of their findings (Cole et al., 2013; Garrison et al., 2015; Hong et al., 2019), this is not the norm. Based on our analysis applying a relatively conservative threshold can help improve the reliability and prediction accuracy of gradients. This is not surprising, as thresholding is intended to remove noisy connections, which contain limited signal. As would be expected, our work also found that highly stringent thresholds result in diminishing returns and eventual loss of signal. As such, threshold selections should ideally be based on exhaustive searches of threshold parameters on test datasets that can be used to assess reliability and validity, prior to application to datasets of interest. Of note, our work also examined the impact of the nature of the thresholding approach (*i.e*., row-wise *vs*. global [applied to full matrix]), finding little differences in these changes.

Building on an emerging theme in the functional connectomics literature, the present work found that “more data is better” – for both reliability and predictive accuracy. Our vertex-wise reliability analyses using intraclass correlation coefficient showed progressive increases as the amount of data available increased from 5-50 minutes. These findings mimic those of edgewise connectivity and areal parcellation in prior studies (Elliott et al. 2019; Noble et al. 2017; Xu et al. 2016; Zuo et al. 2019; Nikolaidis et al. 2020). Consistent with these works, vertices within the frontoparietal, default and dorsal attention networks showed higher reliability with lower amounts of data. This finding raises the question as to whether the prominence of findings in these networks in the literature may be a reflection of their reliability. Our connectome-wide reliability analyses based on discriminability also supported the value of greater amounts of data, though patterns of change were less progressive, advantages were only found for going from 5 to 10 minutes of data per subject, and then from 30 to 50 minutes. As would be expected based on the increases in reliability observed in both univariate and multivariate perspectives, the significance of predictive relationships also increased with data availability, regardless of the domain of behavior being examined. Of note, from multivariate perspectives, compared to the ICC, discriminability may provide a more explicit criterion to determine a predictive power of gradients. Indeed, as shown in **Figure 4**, nearly 1 value of discriminability usually confirmed FDR-significant phenotypic predictions, whereas whole-brain averaged ICC revealed significant prediction across even the ranges that are considered as “poor” and somewhat arbitrary (=0.2-0.3).

A key limitation of the present work is that the connectivity gradients framework was notably more successful in predicting cognitive than psychiatric variables. We do not believe this should be taken to infer that connectivity gradients are not relevant to psychopathology. Instead, we posit that the lower performance with psychiatric variables is likely a reflection of the composition of the Human Connectome Project, which was largely focused on neurotypical adults. In such a population, the magnitude of differences among individuals would be expected to be smaller. Compounding this expectation, is the reality that questionnaire instruments such as the Adult Self Report are largely designed to differentiate individuals based on the severity of psychiatric symptoms, which makes them relatively limited in the assessment of differences among neurotypical adults. One other limitation to note is that while we demonstrated the feasibility of improving prediction accuracy by combining multiple gradients, our effort to identify those gradients most likely to be contribute meaningfully was found to be suboptimal for externalizing symptoms. While this may have actually reflected limitations in the composition or size of the training set, further examination of methods for combining gradients to optimize prediction is clearly merited.

Finally, it is worth noting the need for a clearer conceptual positioning of connectivity gradients with respect to conventional network detection approaches. The networks, which are commonly revealed by independent component analysis (ICA) or clustering techniques, have constituted a dominant unit of brain organization until recently. Both approaches aim to describe the same highdimensional brain connectivity space, yet with different perspectives. For example, while ICA-based functional networks can be conceptualized as spatially distinct sources within the connectivity matrix, PCA-derived gradients show rather smoother transition of brain areas along the continuous dimension capturing topographic connectivity relationships. Mathematically, PCA finds multiple orthogonal directions that can explain maximal variance of Gaussian data, whereas ICA seeks independent axes which direction indicates the ‘sources’ of original non-Gaussian data. Because of such conceptual and mathematical differences between the two methods, the spatial correlation of their components (PCA-derived gradients *vs*. non-thresholded ICA maps) are only moderate (0.3-0.5 on average), even if they are extracted from the same subject, compared to those between PCA, DE and LE (**Supplementary Figure 6**). The challenge moving forward is in the judicious application of these interrelated but distinct perspectives to capturing the relevant features of brain organization; in the end, it may be the combination of both perspectives that is needed to provide more comprehensive solutions for biomarker discovery.

### Recommendations

The results of the present work serve as a starting point for those interested in pursuing biomarker discovery using connectivity gradients. From this perspective, in the absence of further testing, we would recommend usage of the following parameters in studies using gradient based approaches:

1. Consider the thresholding of a functional connectivity matrix.
2. Limit analysis to the most reliable gradients.
3. ICC may be arbitrary for selecting gradients; discriminability is more determined and thus preferable.
4. More data the better for voxel-wise ICC and whole-brain discriminability, and the standard 5-10 minutes are suboptimal.

### Conclusions

The present work aimed to establish a set of benchmark parameters for the use of low dimensional representation of functional connectivity in the domain of biomarker discovery. We evaluated parameters to maximize reliability of this approach while retaining its ability to predict individuals. Our results highlight the importance of carefully considering the choice of algorithms, the degree of thresholding and the length of scan since each of these choices impacts on the ability of gradient representations of the cortex as a tool for biomarker discovery. We hope that these findings will help provide a foundation with which researchers can efficiently use gradient framework in their research. Finally, it is worth noting that our findings emphasize the importance of considering the amount of data employed for the calculation of features and its implications for reliability — the latter of which is a primary determinant of sample size needs for detection of targeted effects, as well as the feasibility of making individual level predictions.

## Acknowledgements

The authors wish to thank all the efforts of the HCP and MSC teams to create and maintain the open sharing data repositories. This work was supported by funding from the Brain & Behavior Research Foundation (NARSAD Young Investigator grant; #28436), the Canadian Institutes of Health Research (postdoctoral fellowship MFE-158228) and the Institute for Basic Science (IBS-R15-D1) for SJH; the National Institute of Mental Health (R21MH118556-02) and the Brain & Behavior Research Foundation (NARSAD Young Investigator grant; #12345) for AN; the European Research Council (WANDERINGMINDS – ERC646927) for JS; National Science and Engineering Research Council of Canada (NSERC, Discovery-1304413), the Canadian Institutes of Health Research (CIHR, FDN-154298), the Azrieli Center for Autism Research of the Montreal Neurological Institute, SickKids Foundation (NI17-039), and Canada Research Chairs Program for BCB; the National Institute of Mental Health (U01MH099059), and an endowment from the Phyllis Green and Randolph Cowen for MPM.

## References

Alexander, L.M., Salum, G.A., Swanson, J.M., Milham, M.P., 2020. Measuring strengths and weaknesses in dimensional psychiatry. Journal of Child Psychology and Psychiatry 61, 40–50.

Barch, D.M., Burgess, G.C., Harms, M.P., Petersen, S.E., Schlaggar, B.L., Corbetta, M., Glasser, M.F., Curtiss, S., Dixit, S., Feldt, C., Nolan, D., Bryant, E., Hartley, T., Footer, O., Bjork, J.M., Poldrack, R., Smith, S., Johansen-Berg, H., Snyder, A.Z., Van Essen, D.C., WU-Minn HCP Consortium, 2013. Function in the human connectome: task-fMRI and individual differences in behavior. Neuroimage 80, 169–189.

Belkin, M., Niyogi, P., 2003. Laplacian Eigenmaps for Dimensionality Reduction and Data Representation. Neural Comput. 15, 1373–1396.

Benjamini, Y., Hochberg, Y., 1995. Controlling the False Discovery Rate: A Practical and Powerful Approach to Multiple Testing. J. R. Stat. Soc. Series B Stat. Methodol. 57, 289–300.

Birn, R.M., Molloy, E.K., Patriat, R., Parker, T., Meier, T.B., Kirk, G.R., Nair, V.A., Meyerand, M.E., Prabhakaran, V., 2013. The effect of scan length on the reliability of resting-state fMRI connectivity estimates. Neuroimage 83, 550–558.

Bridgeford, E.W., Wang, S., Yang, Z., Wang, Z., Xu, T., Craddock, C., Dey, J., Kiar, G., Gray-Roncal, W., Priebe, C.E., Caffo, B., Milham, M., Zuo, X.-N., Consortium for Reliability and Reproduciblity, Vogelstein, J.T., 2020. Big Data Reproducibility: Applications in Brain Imaging. bioRxiv. https://doi.org/10.1101/802629

Bullmore, E., Sporns, O., 2009. Complex brain networks: graph theoretical analysis of structural and functional systems. Nat. Rev. Neurosci. 10, 186–198.

Castellanos, F.X., Xavier Castellanos, F., Di Martino, A., Cameron Craddock, R., Mehta, A.D., Milham, M.P., 2013. Clinical applications of the functional connectome. NeuroImage. https://doi.org/10.1016/j.neuroimage.2013.04.083

Catani, M., Ffytche, D.H., 2005. The rises and falls of disconnection syndromes. Brain 128, 2224–2239.

Cohen, A.L., Fair, D.A., Dosenbach, N.U.F., Miezin, F.M., Dierker, D., Van Essen, D.C., Schlaggar, B.L., Petersen, S.E., 2008. Defining functional areas in individual human brains using resting functional connectivity MRI. Neuroimage 41, 45–57.

Coifman, R.R., Lafon, S., 2006. Diffusion maps. Appl. Comput. Harmon. Anal. 21, 5–30.

Coifman, R.R., Lafon, S., Lee, A.B., Maggioni, M., Nadler, B., Warner, F., Zucker, S.W., 2005. Geometric diffusions as a tool for harmonic analysis and structure definition of data: diffusion maps. Proc. Natl. Acad. Sci. U. S. A. 102, 7426–7431.

Cole, M.W., Reynolds, J.R., Power, J.D., Repovs, G., Anticevic, A., Braver, T.S., 2013. Multitask connectivity reveals flexible hubs for adaptive task control. Nat. Neurosci. 16, 1348–1355.

Craddock, R.C., James, G.A., Holtzheimer, P.E., 3rd, Hu, X.P., Mayberg, H.S., 2012. A whole brain fMRI atlas generated via spatially constrained spectral clustering. Hum. Brain Mapp. 33, 1914–1928.

Di Martino, A., Fair, D.A., Kelly, C., Satterthwaite, T.D., Castellanos, F.X., Thomason, M.E., Craddock, R.C., Luna, B., Leventhal, B.L., Zuo, X.-N., Milham, M.P., 2014. Unraveling the miswired connectome: a developmental perspective. Neuron 83, 1335–1353.

Duncan, J., 2010. The multiple-demand (MD) system of the primate brain: mental programs for intelligent behaviour. Trends Cogn. Sci. 14, 172–179.

Elliott, M.L., Knodt, A.R., Cooke, M., Kim, M.J., Melzer, T.R., Keenan, R., Ireland, D., Ramrakha, S., Poulton, R., Caspi, A., Moffitt, T.E., Hariri, A.R., 2019. General functional connectivity: Shared features of resting-state and task fMRI drive reliable and heritable individual differences in functional brain networks. Neuroimage 189, 516–532.

Fox, M.D., Snyder, A.Z., Vincent, J.L., Corbetta, M., Van Essen, D.C., Raichle, M.E., 2005. The human brain is intrinsically organized into dynamic, anticorrelated functional networks. Proc. Natl. Acad. Sci. U. S. A. 102, 9673–9678.

Garrison, K.A., Scheinost, D., Finn, E.S., Shen, X., Constable, R.T., 2015. The (in)stability of functional brain network measures across thresholds. Neuroimage 118, 651–661.

Garrity, A.G., Pearlson, G.D., McKiernan, K., Lloyd, D., Kiehl, K.A., Calhoun, V.D., 2007. Aberrant “Default Mode” Functional Connectivity in Schizophrenia. AJP 164, 450–457.

Glasser, M.F., Sotiropoulos, S.N., Wilson, J.A., Coalson, T.S., Fischl, B., Andersson, J.L., Xu, J., Jbabdi, S., Webster, M., Polimeni, J.R., Van Essen, D.C., Jenkinson, M., WU-Minn HCP Consortium, 2013. The minimal preprocessing pipelines for the Human Connectome Project. Neuroimage 80, 105–124.

Gordon, E.M., Laumann, T.O., Gilmore, A.W., Newbold, D.J., Greene, D.J., Berg, J.J., Ortega, M., Hoyt-Drazen, C., Gratton, C., Sun, H., Hampton, J.M., Coalson, R.S., Nguyen, A.L., McDermott, K.B., Shimony, J.S., Snyder, A.Z., Schlaggar, B.L., Petersen, S.E., Nelson, S.M., Dosenbach, N.U.F., 2017. Precision Functional Mapping of Individual Human Brains. Neuron. https://doi.org/10.1016/j.neuron.2017.07.011

Greicius, M.D., Flores, B.H., Menon, V., Glover, G.H., Solvason, H.B., Kenna, H., Reiss, A.L., Schatzberg, A.F., 2007. Resting-state functional connectivity in major depression: abnormally increased contributions from subgenual cingulate cortex and thalamus. Biol. Psychiatry 62, 429–437.

Haak, K.V., Marquand, A.F., Beckmann, C.F., 2018. Connectopic mapping with resting-state fMRI. Neuroimage 170, 83–94.

Hilgetag, C.C., Goulas, A., 2020. “Hierarchy” in the organization of brain networks. Philos. Trans. R. Soc. Lond. B Biol. Sci. 375, 20190319.

Hong, S.-J., Vos de Wael, R., Bethlehem, R.A.I., Lariviere, S., Paquola, C., Valk, S.L., Milham, M.P., Di Martino, A., Margulies, D.S., Smallwood, J., Bernhardt, B.C., 2019. Atypical functional connectome hierarchy in autism. Nat. Commun. 10, 1022.

Landis, J.R., Koch, G.G., 1977. The measurement of observer agreement for categorical data. Biometrics 33, 159–174.

Langs, G., Sweet, A., Lashkari, D., Tie, Y., Rigolo, L., Golby, A.J., Golland, P., 2014. Decoupling function and anatomy in atlases of functional connectivity patterns: language mapping in tumor patients. Neuroimage 103, 462–475.

Langs, G., Wang, D., Golland, P., Mueller, S., Pan, R., Sabuncu, M.R., Sun, W., Li, K., Liu, H., 2016. Identifying Shared Brain Networks in Individuals by Decoupling Functional and Anatomical Variability. Cereb. Cortex 26, 4004–4014.

Margulies, D.S., Ghosh, S.S., Goulas, A., Falkiewicz, M., Huntenburg, J.M., Langs, G., Bezgin, G., Eickhoff, S.B., Castellanos, F.X., Petrides, M., Jefferies, E., Smallwood, J., 2016. Situating the default-mode network along a principal gradient of macroscale cortical organization. Proc. Natl. Acad. Sci. U. S. A. 113, 12574–12579.

Marquand, A.F., Haak, K.V., Beckmann, C.F., 2017. Functional corticostriatal connection topographies predict goal directed behaviour in humans. Nat Hum Behav 1, 0146.

Mars, R.B., Passingham, R.E., Jbabdi, S., 2018a. Connectivity Fingerprints: From Areal Descriptions to Abstract Spaces. Trends in Cognitive Sciences. https://doi.org/10.1016/j.tics.2018.08.009

Mars, R.B., Sotiropoulos, S.N., Passingham, R.E., Sallet, J., Verhagen, L., Khrapitchev, A.A., Sibson, N., Jbabdi, S., 2018b. Whole brain comparative anatomy using connectivity blueprints. Elife 7. https://doi.org/10.7554/eLife.35237

McIntosh, A.R., Mišić, B., 2013. Multivariate statistical analyses for neuroimaging data. Annu. Rev. Psychol. 64, 499–525.

Menon, V., 2011. Large-scale brain networks and psychopathology: a unifying triple network model. Trends Cogn. Sci. 15, 483–506.

Mesulam, M.M., 1998. From sensation to cognition. Brain 121 (Pt 6), 1013–1052.

Murphy, C., Jefferies, E., Rueschemeyer, S.-A., Sormaz, M., Wang, H.-T., Margulies, D.S., Smallwood, J., 2018. Distant from input: Evidence of regions within the default mode network supporting perceptually-decoupled and conceptually-guided cognition. Neuroimage 171, 393–401.

Murphy, C., Wang, H.-T., Konu, D., Lowndes, R., Margulies, D.S., Jefferies, E., Smallwood, J., 2019. Modes of operation: A topographic neural gradient supporting stimulus dependent and independent cognition. Neuroimage 186, 487–496.

Navarro Schröder, T., Haak, K.V., Zaragoza Jimenez, N.I., Beckmann, C.F., Doeller, C.F., 2015. Functional topography of the human entorhinal cortex. Elife 4. https://doi.org/10.7554/eLife.06738

Nikolaidis, A., Heinsfeld, A.S., Xu, T., Bellec, P., Vogelstein, J., Milham, M., 2020. Bagging improves reproducibility of functional parcellation of the human brain. Neuroimage 116678.

Noble, S., Spann, M.N., Tokoglu, F., Shen, X., Constable, R.T., Scheinost, D., 2017. Influences on the Test-Retest Reliability of Functional Connectivity MRI and its Relationship with Behavioral Utility. Cereb. Cortex 27, 5415–5429.

Jolliffe, I., 2011. Principal Component Analysis. In: Lovric, M. (Ed.), International Encyclopedia of Statistical Science. Springer Berlin Heidelberg, Berlin, Heidelberg, pp. 1094–1096.

Przeździk, I., Faber, M., Fernández, G., Beckmann, C.F., Haak, K.V., 2019. The functional organisation of the hippocampus along its long axis is gradual and predicts recollection. Cortex 119, 324–335.

Roalf, D.R., Gur, R.C., 2017. Functional brain imaging in neuropsychology over the past 25 years. Neuropsychology 31, 954–971.

Salimi-Khorshidi, G., Douaud, G., Beckmann, C.F., Glasser, M.F., Griffanti, L., Smith, S.M., 2014. Automatic denoising of functional MRI data: combining independent component analysis and hierarchical fusion of classifiers. Neuroimage 90, 449–468.

Schaefer, A., Kong, R., Gordon, E.M., Laumann, T.O., Zuo, X.-N., Holmes, A.J., Eickhoff, S.B., Yeo, B.T.T., 2018. Local-Global Parcellation of the Human Cerebral Cortex from Intrinsic Functional Connectivity MRI. Cereb. Cortex 28, 3095–3114.

Shen, X., Finn, E.S., Scheinost, D., Rosenberg, M.D., Chun, M.M., Papademetris, X., Constable, R.T., 2017. Using connectome-based predictive modeling to predict individual behavior from brain connectivity. Nat. Protoc. 12, 506–518.

Shrout, P.E., Fleiss, J.L., 1979. Intraclass correlations: uses in assessing rater reliability. Psychol. Bull. 86, 420–428.

Smith, S.M., Beckmann, C.F., Andersson, J., Auerbach, E.J., Bijsterbosch, J., Douaud, G., Duff, E., Feinberg, D.A., Griffanti, L., Harms, M.P., Kelly, M., Laumann, T., Miller, K.L., Moeller, S., Petersen, S., Power, J., Salimi-Khorshidi, G., Snyder, A.Z., Vu, A.T., Woolrich, M.W., Xu, J., Yacoub, E., Uğurbil, K., Van Essen, D.C., Glasser, M.F., WU-Minn HCP Consortium, 2013. Resting-state fMRI in the Human Connectome Project. Neuroimage 80, 144–168.

Smith, S.M., Nichols, T.E., Vidaurre, D., Winkler, A.M., Behrens, T.E.J., Glasser, M.F., Ugurbil, K., Barch, D.M., Van Essen, D.C., Miller, K.L., 2015. A positive-negative mode of population covariation links brain connectivity, demographics and behavior. Nat. Neurosci. 18, 1565–1567.

Sormaz, M., Murphy, C., Wang, H.-T., Hymers, M., Karapanagiotidis, T., Poerio, G., Margulies, D.S., Jefferies, E., Smallwood, J., 2018. Default mode network can support the level of detail in experience during active task states. Proc. Natl. Acad. Sci. U. S. A. 115, 9318–9323.

van den Heuvel, M.P., Sporns, O., 2019. A cross-disorder connectome landscape of brain dysconnectivity. Nat. Rev. Neurosci. 20, 435–446.

Van Den Heuvel, M.P., Sporns, O., 2011. Rich-club organization of the human connectome. Journal of Neuroscience.

Vos de Wael, R., Benkarim, O., Paquola, C., Lariviere, S., Royer, J., Tavakol, S., Xu, T., Hong, S.-J., Langs, G., Valk, S., Misic, B., Milham, M., Margulies, D., Smallwood, J., Bernhardt, B.C., 2020. BrainSpace: a toolbox for the analysis of macroscale gradients in neuroimaging and connectomics datasets. Commun Biol 3, 103.

Vos de Wael, R., Larivière, S., Caldairou, B., Hong, S.-J., Margulies, D.S., Jefferies, E., Bernasconi, A., Smallwood, J., Bernasconi, N., Bernhardt, B.C., 2018. Anatomical and microstructural determinants of hippocampal subfield functional connectome embedding. Proc. Natl. Acad. Sci. U. S. A. 115, 10154–10159.

Wang, C., Mahadevan, S., 2008. Manifold alignment using Procrustes analysis, in: Proceedings of the 25th International Conference on Machine Learning, ICML ’08. Association for Computing Machinery, New York, NY, USA, pp. 1120–1127.

Wang, H.-T., Smallwood, J., Mourao-Miranda, J., Xia, C.H., Satterthwaite, T.D., Bassett, D.S., Bzdok, D., 2018. Finding the needle in high-dimensional haystack: A tutorial on canonical correlation analysis. arXiv [stat.ML].

Wig, G.S., Laumann, T.O., Petersen, S.E., 2014. An approach for parcellating human cortical areas using resting-state correlations. Neuroimage 93 Pt 2, 276–291.

Xu, T., Opitz, A., Craddock, R.C., Wright, M.J., Zuo, X.-N., Milham, M.P., 2016. Assessing Variations in Areal Organization for the Intrinsic Brain: From Fingerprints to Reliability. Cereb. Cortex 26, 4192–4211.

Xu, T., Yang, Z., Jiang, L., Xing, X.-X., Zuo, X.-N., 2015. A Connectome Computation System for discovery science of brain. Sci Bull. Fac. Agric. Kyushu Univ. 60, 86–95.

Yeo, B.T.T., Krienen, F.M., Sepulcre, J., Sabuncu, M.R., Lashkari, D., Hollinshead, M., Roffman, J.L., Smoller, J.W., Zöllei, L., Polimeni, J.R., Fischl, B., Liu, H., Buckner, R.L., 2011. The organization of the human cerebral cortex estimated by intrinsic functional connectivity. J. Neurophysiol. 106, 1125–1165.

Zuo, X.-N., Xing, X.-X., 2014. Test-retest reliabilities of resting-state FMRI measurements in human brain functional connectomics: a systems neuroscience perspective. Neurosci. Biobehav. Rev. 45, 100–118.

Zuo, X.-N., Xu, T., Milham, M.P., 2019. Harnessing reliability for neuroscience research. Nat Hum Behav 3, 768–771.

