## Supplementary materials for "TOWARD A CONNECTIVITY GRADIENT-BASED FRAMEWORK FOR REPRODUCIBLE BIOMARKER DISCOVERY"

**Hong SJ, Xu T, et al.**

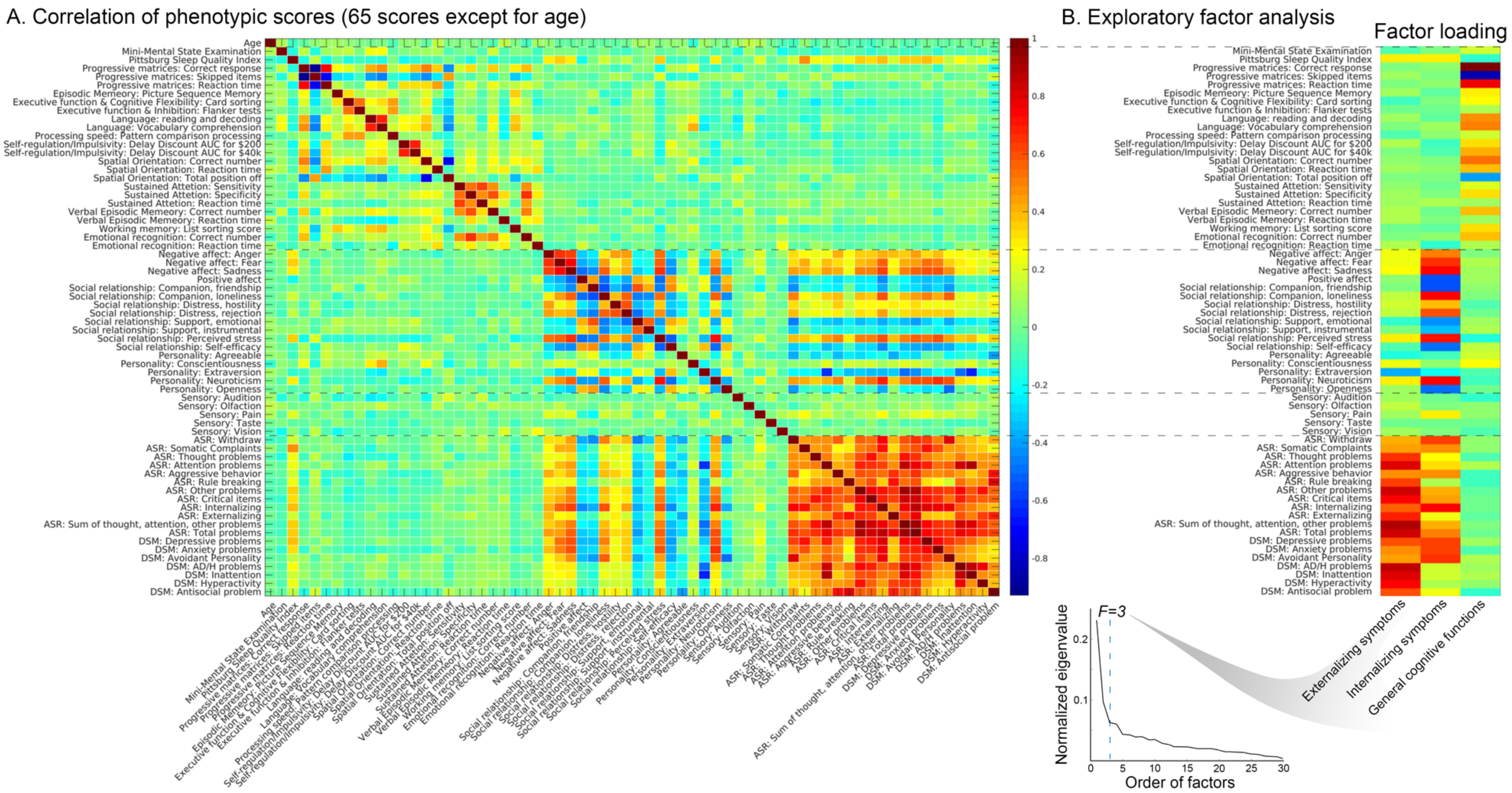


**Supplementary Figure 1. Phenotypic score correlation and exploratory factor analysis. A**) The correlation matrix across 65 phenotypic scores + age is shown. **B**) The three factor models (*i.e.,* externalizing, internalizing and general cognitive profiles) derived from an exploratory factor analysis is present, together with a list of phenotypic scores consisting of each factor next. The optimal model was chosen at the order showing an elbow point (=3) based on a normalized eigenvalue.

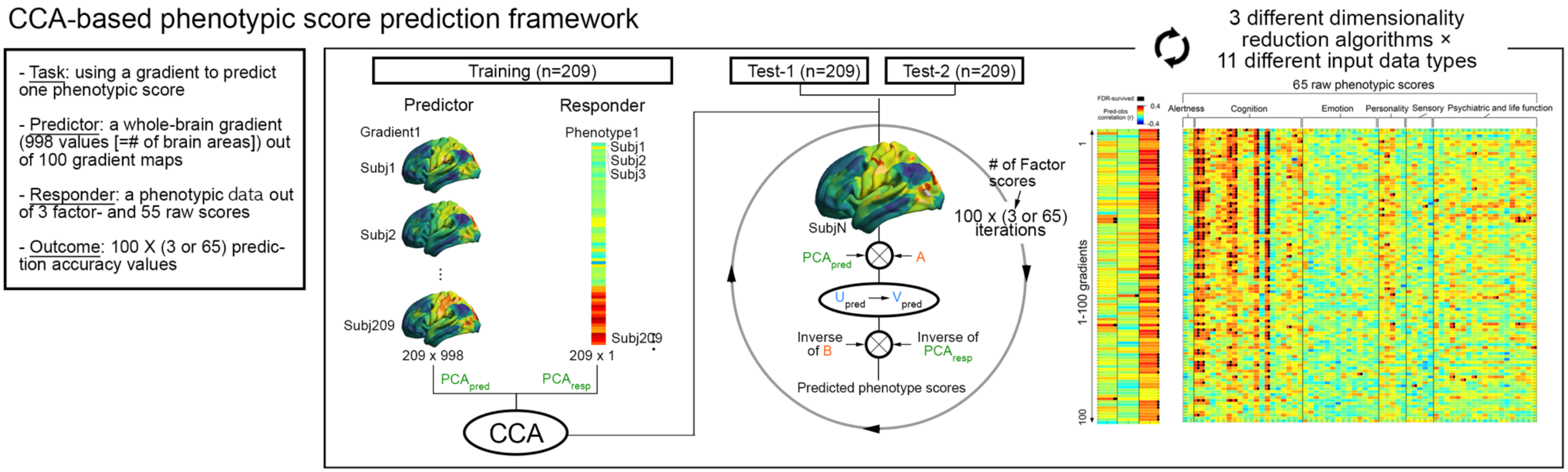


**Supplementary Figure 2. Methodological details for our canonical correlation analysis (CCA)-based prediction framework.** The main goal of this prediction was to predict a single phenotypic score (a responder: 209×1) based on a single gradient map (a predictor: 209×998 [# of parcels]). For generalizability, we trained our framework using a discovery cohort (a training dataset), while testing based on completely independent two datasets (test-1 and -2). The outcomes were 100 × 3/65 prediction accuracy tables, where 100 refers to the number of targeted gradients, 3 to the number of factor scores and 65 to the number of the entire phenotypic scores. The core of the prediction framework was CCA, by which we sought to find an optimal linear relationship between gradient values and a phenotypic score in the training dataset. The entire framework consisted of 5 steps: *i)* performing PCA on the input gradient of a training dataset to reduce its dimensionality (209×998 to 209×Y), *ii)* performing CCA between reduced gradient maps and a phenotypic scores to obtain canonical coefficients (A, B), *iii)* applying the *learned* PCA coefficients to the gradient maps of test cases, *iv)* applying the *learned* canonical coefficients (A) to the reduced gradient maps of test cases, *v)* inverse mapping of PCA and CCA coefficients (B) to the outcome of the previous step (*iv*), which provides reconstructed individual phenotypic scores as a prediction result. This entire framework was repeatedly performed 3×11 times to test different combinations between three dimensionality reduction algorithms and 11 input data types. See '*2) Prediction framework*' in *Analysis-4* of **METHODS** for details.

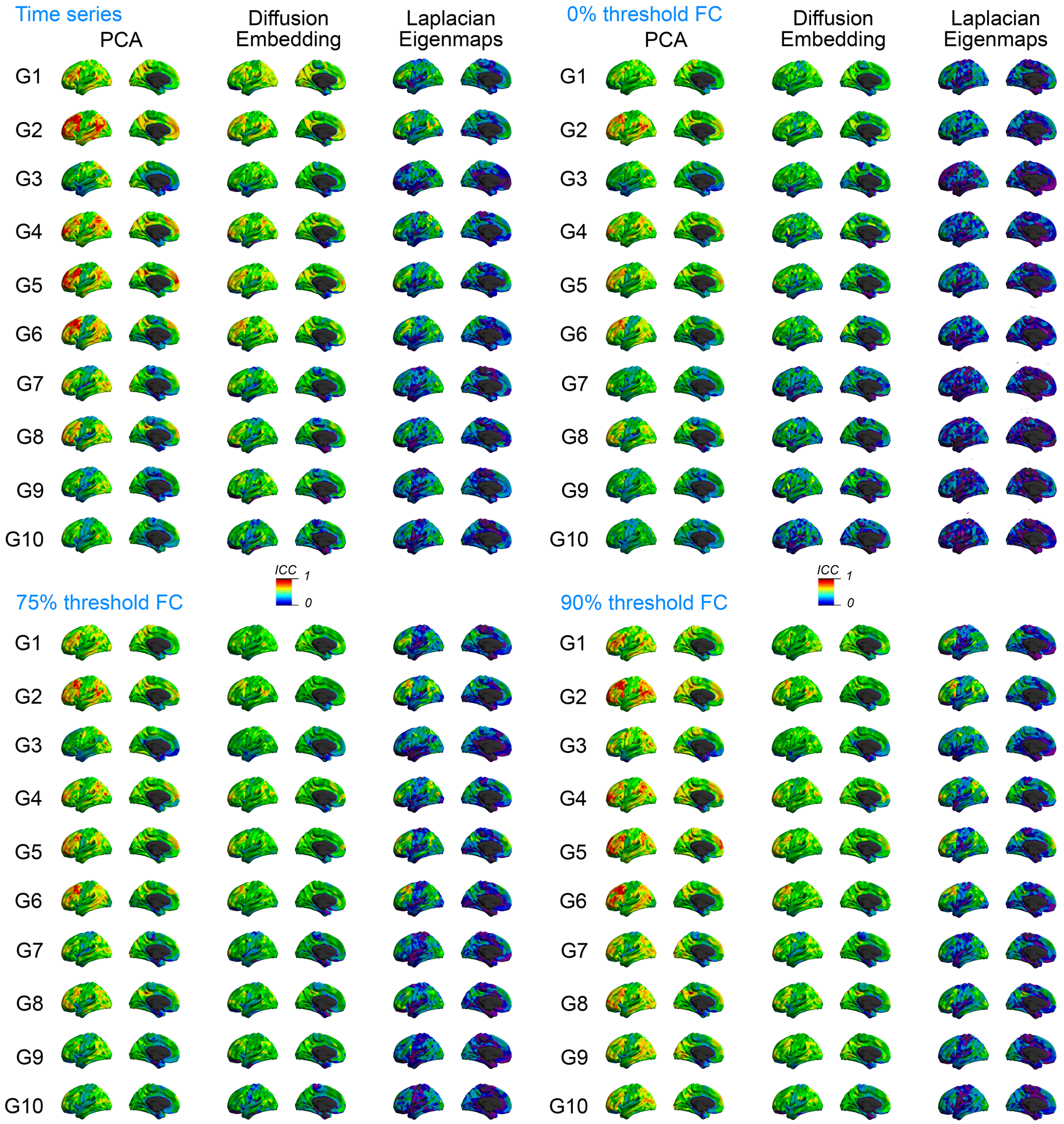


**Supplementary Figure 3. Reliability maps of gradients across different combinations of gradient calculation parameters.** Intraclass correlation coefficients of functional gradients were mapped on the whole brain across different combinations of algorithms (*i.e.,* PCA, diffusion embedding and Laplacian Eigenmaps) and input data types (*i.e.,* time series, 0%, 75% and 90% thresholded functional connectivity matrices). The general spatial patterns across the algorithms are highly reproducible.

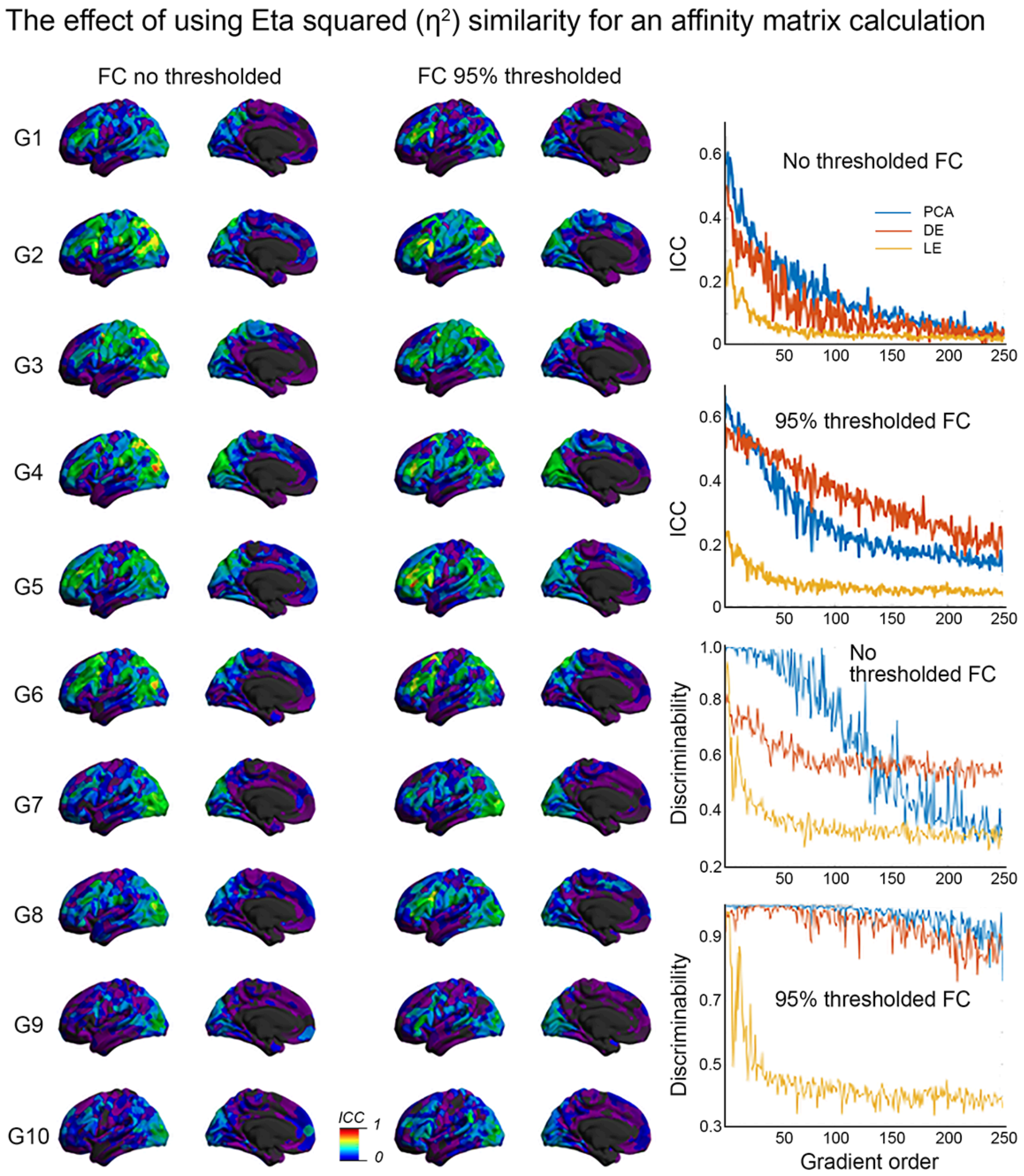


**Supplementary Figure 4. The gradient and reliability profiles of Laplacian Eigenmaps based on an affinity matrix derived from Eta-squared ($\eta^2$) similarity.** To test the effect of a different approach for affinity matrix calculation on reliability, we re-generated ICC maps of the gradients derived from a $\eta^2$ similarity kernel, following the previous study (Haak et al., 2018). (*Left*) As examples, we chose 10 gradients extracted from 0% and 95% thresholded functional connectivity matrices. (*Right*) we also presented the whole brain ICC and discriminability changes as a function of gradient order, comparing their patterns between the different algorithms (PCA, diffusion embedding and Laplacian Eigenmaps).

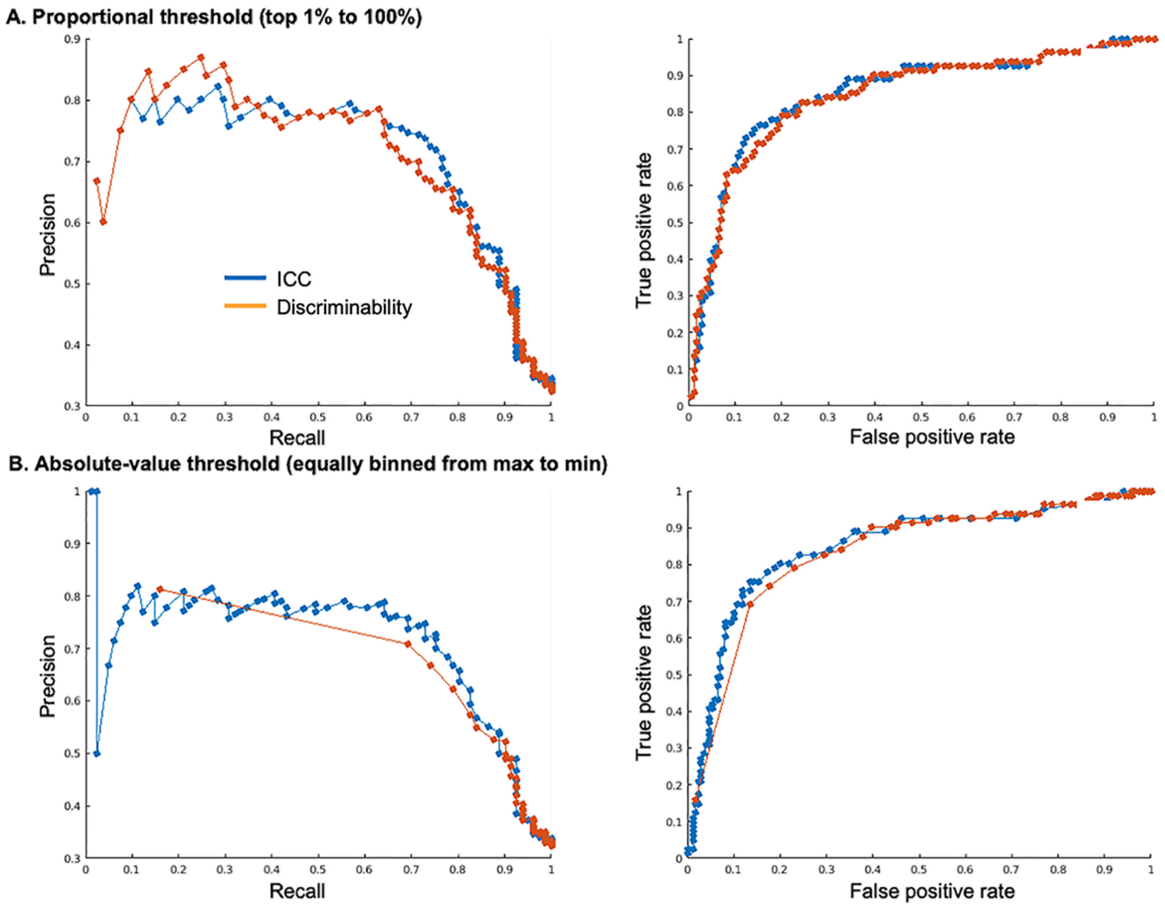


**Supplementary Figure 5. Precision-Recall and Receiver-Operating Characteristic curves to compare the univariate ICC and multivariate discriminability.** The aim of this analysis was to assess which one between the two reliability measures (*i.e.,* ICC *vs.* discriminability) more accurately infers the significant phenotypic prediction. To this end, we constructed two metrics for evaluating a prediction result, namely precision-recall and receiver-operating characteristic (ROC) curves. For simplification, here we used prediction results for three factor scores (instead of 65-phenotypic-scores based results). **A**) In the first experiment, we *i)* systematically decreased each reliability measure from top 1% to 100% (every 1% interval) to define a threshold, *ii)* each time, aggregated the gradients with ICC greater than the defined threshold, and *iii)* assessed how many those gradients were able to predict at least one factor scores or not. By doing this, we could construct precision, recall and false positive rate values across different reliability thresholds, which allowed for constructing precision-recall (*left*) and ROC (*right*) curves. As shown in the graphs, the two reliability measures do not show noticeable differences in both evaluation curves. **B**) Conceptually same analyses were repeated but with an alternative thresholding strategy (systematically decreasing absolute value of reliability measures). Indeed, this time, we decreased a small amount of the threshold from the maximum reliability value to zero iteratively and assessed if those gradients with ICC>the threshold can predict phenotypic scores or not. Again, we constructed precision-recall and ROC curves from this analysis and found that they do not necessarily differ in terms of inference ability for significant phenotypic prediction. Nevertheless, one noticeable finding was that in discriminability, a very high threshold (nearly perfect discriminability) already showed a certain level of inference ability, suggesting its sensitivity compared to ICC.

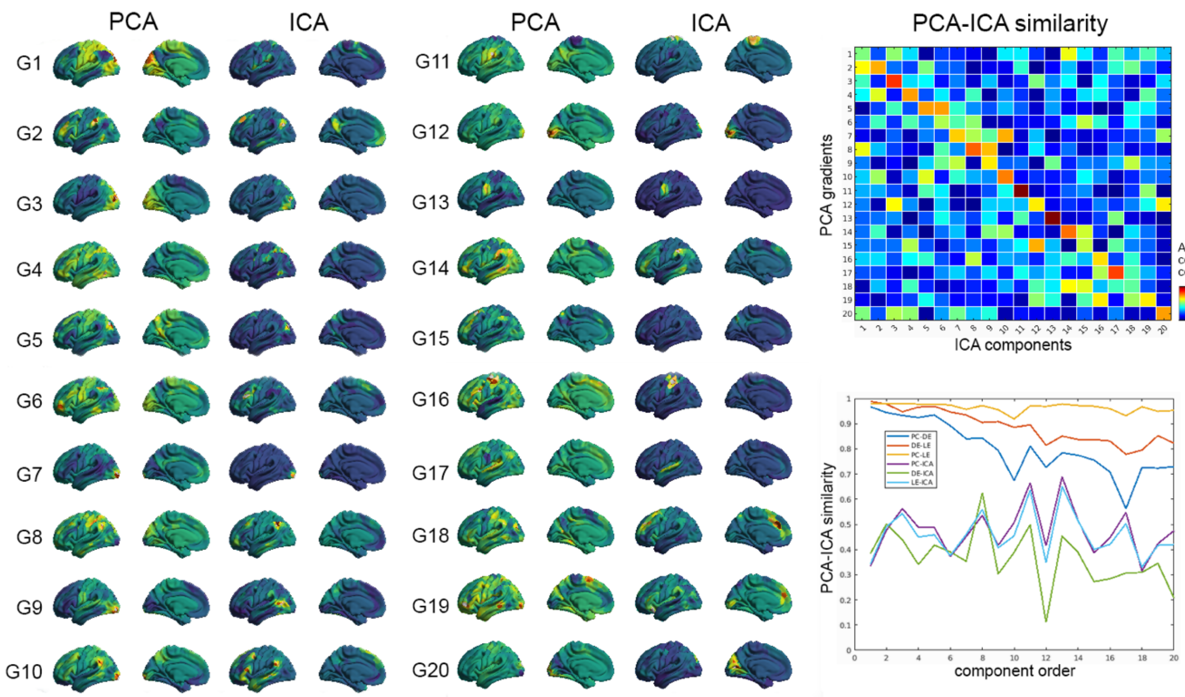


**Supplementary Figure 6. PCA and ICA component profiles and their (di)similarity.** (*Left*) 20 primary components from PCA and ICA are mapped on the whole-brain cortex. Using a Hungarian matching, the component orders are matched (but not for the direction of sign for transparency). (*Right, top*) Across all components, spatial similarity (absolute correlation coefficient) was computed across a pair of gradient orders. Note that after Hungarian matching, the values along the diagonal elements (*i.e.,* similarity for the same order gradient) became improved. (*Right, bottom*) The similarity of ICA towards all three dimensionality reduction algorithms analyzed in our study was presented along the component order. Compared to within-dimensionality reduction algorithms, ICA shows generally lower similarity, regardless of the order.
